# Dynamic encoding of saccade sequences in primate frontal eye field

**DOI:** 10.1101/2020.08.01.232488

**Authors:** Jing Jia, Zhen Puyang, Qingjun Wang, Xin Jin, Aihua Chen

## Abstract

Frontal eye field (FEF) is a key part of oculomotor system, with dominant responses to the direction of single saccades. However, whether and how FEF contributes to sequential saccades remain largely unknown. Here by training rhesus monkeys to perform sequential saccades and recording the neuronal activities in FEF, we found that the sequence-related activities are clearly represented in FEF, and many neurons’ selectivity to saccade direction undergoes dynamic changes during sequential task. In addition, the sequence-related activities are context-dependent, with different firing activities during memory- versus visually-guided sequence. Supra-threshold microstimulation in FEF evokes saccade without altering the overall sequence structure. Pharmacological inactivation of FEF severely impaired the monkey’s performance of sequential saccades, with different effects on the same actions at different positions within the sequence. These results reveal the context-dependent dynamic encoding of saccade direction in FEF, and underscore a critical role of FEF in planning and execution of sequential saccades.

**In Brief:** Jia, Puyang et al. employed in vivo recording to reveal the dynamic encoding of sequential saccades in primate frontal eye field (FEF), then used electric microstimulation and reversible inactivation to demonstrate the causal role of FEF in controlling saccade sequences.

**Highlights:**

- FEF neurons respond differently during sequential vs. single saccades
- Sequence-related FEF activity is context-dependent
- FEF microstimulation induced saccade without altering sequence structure
- FEF inactivation severely impaired the performance of sequential saccades

## Introduction

The primate frontal eye field (FEF) is known as a critical area for saccade movement (Schall JD 2002). Lots of neurophysiological studies carried in FEF demonstrated that it plays an important role in transforming the outside visual signals into a command of saccade to certain direction (Schall JD 2004). Accordingly, previous study on the orderly arrangement of three center-out saccades in FEF indicated that it was largely involved in the direction selection of forthcoming saccades, but barely in the temporal structure of sequence (Isoda M and J Tanji 2003). While during the oculomotor performance each saccade has to follow one another within a sequence, accumulating evidences have suggested that the organization of sequence might not be serial but hierarchical (Jin X et al. 2014; Geddes CE et al. 2018), and the planning of saccade sequence could be parallel (McPeek RM et al. 2000; Isoda M and J Tanji 2003; Schall JD 2004; Basu D and A Murthy 2020). Through this way, it can lead to effective execution of a whole sequence at the same time with certain tolerance of error behavior (Jin X *et al*. 2014; Geddes CE *et al*. 2018). Indeed, it has been previously reported that the neurons in supplementary eye field demonstrate context-dependent selectivity of saccadic direction and may contribute to the planning and generation of learned oculomotor sequences (Lu X et al. 2002; Isoda M and J Tanji 2003). These findings thus raise the questions of wheer and how FEF encodes and causally contributes to sequential saccades.

To investigate these questions, we trained rhesus macaques to perform a novel self-paced oculomotor sequence task consisting of four (Left-Left-Right-Right) consecutive saccades, which allow a good internal control for studying the sequence vs. direction-related activities (Geddes CE *et al*. 2018). By comparing the same FEF neuron’s activities during sequential saccades and traditional single memory saccade task, and we found that FEF uses a dynamically encoding strategy for sequential saccade task, and some FEF neurons show activities related with the initiation, switch and termination of the sequence. In addition, these sequence-related activities are context-dependent, with different response characteristics for visual- vs. memory-guided saccade sequences.

To further investigate the dynamic coding characteristics and the causal role of FEF during sequential saccades, electric microstimulation and reversible inactivation in FEF during the performance of sequential saccades were applied, and the results turned out that the supra-threshold microstimulation in FEF evokes saccade but does not affect the overall sequence structure, and interestingly the same direction of saccade at different positions within the sequence was affected differently, indicating that the effect of microstimulation-triggered saccades is more complicated than just generating a saccade to certain direction. Reversible inactivation of FEF severely impaired the performance of sequence, also with the effect varies for the different actions within sequence, rather than only affecting the saccade movement along certain direction. These results indicated that FEF is more complicated than simply generating a vector of saccade movement, and might play an important role for both planning and execution of sequential saccades via dynamic encoding.

## Materials and Methods

### KEY RESOURCES TABLE

| REAGENT or RESOURCE | SOURCE | IDENTIFIER |
| --- | --- | --- |
| Experimental Models:<br>Organisms/Strains |  |  |
| Rhesus Macaque<br>(Macaca Mulatta) | Primate Center | N/A |
| Software and<br>Algorithms |  |  |
| MATLAB | MathWorks | https://www.mathworks.com |
| AlphaLab SnR | AlphaOmega | https://www.aowinery.com/ |
| Spike2 | CED | http://ced.co.uk |
| Tempo | Reflective<br>Computing | http://www.reflectivecomputing.com |
| Other |  |  |
| Tungsten Electrodes | FHC | https://www.fh-co.com |
| Magnetic Field<br>Generator | Crist Instrument | http://www.cristinstrument.com |
| Muscimol | Sigma | https://www.sigmaaldrich.com |

### EXPERIMENTAL MODEL AND SUBJECT DETAILS

In the current study, we used 2 male rhesus monkeys (Macaca mulatta), C and I, weighing 6–8 kg. For each monkey, a lightweight acrylic cap was implanted for head stabilization chronically. Eye movements was monitored through a scleral search coil (Crist Instrument Company, MD, USA). Surgical procedures were performed using aseptic apparatuses under isoflurane anesthesia. After recovery from surgery, animals were trained using standard operant conditioning procedures to perform behavioral tasks to obtain liquid rewards (customized reward system). All animal surgical and experimental procedures were approved by the Institutional Animal Care and Use Committee (IACUC) at East China Normal University.

#### Behavioral Tasks

The experimental setup was similar as previously described (Luna B et al. 1998; Chen A et al. 2016; Shao M et al. 2018). Briefly, monkeys were seated in a primate chair, facing a monitor 57 cm in front of them. A real-time experimental data acquisition (Tempo, Reflective computing, Olympia, WA, USA) and a custum visual stimulus generation system were used to create the behavioral paradigms and record eye position and electrophysiological data. All experiments were conducted in low-level background illumination.

##### Sequential Saccade Task

Monkeys were trained to perform four center-out saccades to the targets in the order of left-left-right-right (L-L-R-R; **Figure 1A**). At the beginning of the task, a white central fixation point (0.3°) and two red target points (0.3°) on the left and right (7° relative to the screen center) were presented. Before sequential saccades, the monkey fixated on a central white fixation point for 100 or 200 ms. Each trial was consisted of two different contexts: visually-guided context and following memory-guided context. For visually-guided sequential task, when the fixation point disappeared, the target point turned white, which indicated the correct saccade direction. If the animal made a saccade to the target within 1500 ms, the target color turned to red again and the central fixation point was illuminated at the same time. Then the animal was required to re-fixate to the central fixation point. The next three saccades were triggered in the same way and correct saccadic sequence was rewarded with a drop of liquid. For memory-guided sequential task, the target points remained red while fixation point disappeared so the monkey needs to remember the sequence of saccades. Animals were also rewarded when finished the memory-guided period. There was no timing restriction for saccades, so monkeys finished this series of saccades at their own pace.

**Figure 1.**
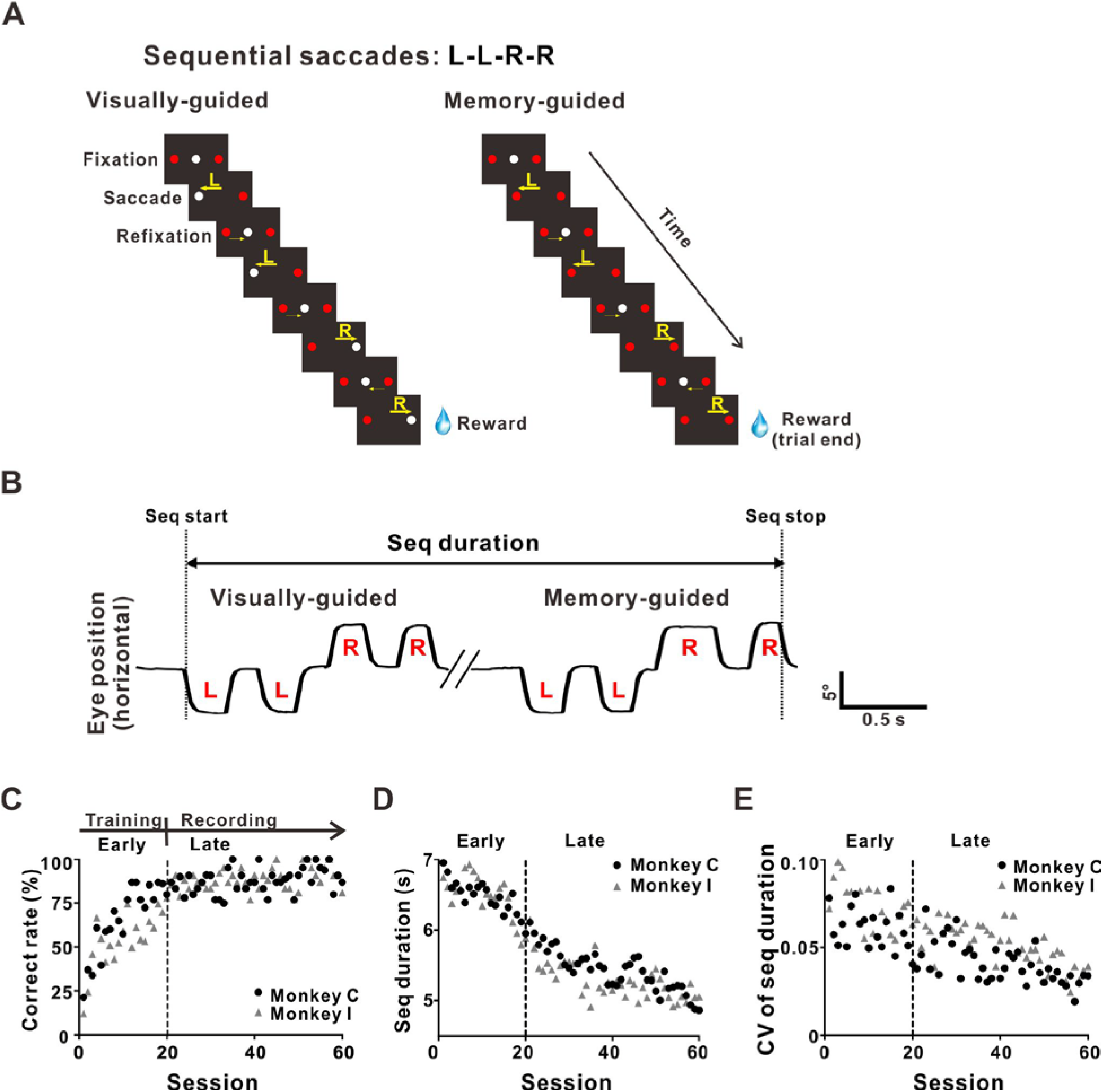
Monkeys learned the sequential saccade task. **(A)** The schematic paradigm of sequence task. Monkeys were trained to perform self-paced visually-guided (left) and memory-guided (right) sequential saccades. In the visually-guided context, saccade direction was cued by the shift of white point (for details, see Method). A white fixation point and two red targets first appeared, and monkey need to make a fixation at the central white dot. For visually-guided sequence, once monkey maintained the fixation for 100 or 200ms, the fixation point disappeared and the target changed the color from red to white indicating that monkey needs to make a saccade to the target within 1500 ms. Once monkey moved to the target, the central fixation point shows up again, and monkey need to make a return saccade to the central refixation point within 1500 ms. Once monkey returned to the central fixation point, the target changed the color again, and monkey need to make a second center-out saccade. In the memory-guided task, there was no cue for saccadic direction to the left or right target. Overall, since saccade is very fast, the limitation of 1500 ms allows that monkey can make saccade sequence according to his own pace for both visually-guided and memory-guided sequence task. One visually-guided sequence was followed by a memory-guide sequence. The sequential order to the saccadic targets was left-left-right-right (L-L-R-R) and the reward was obtained after completing the sequence. **(B)** Samples of a single subject’s eye trace for a full sequence. The x-axis and the y-axis are time and horizontal eye position, respectively. The downward eye trace indicates that monkey moving to the left, while the upward eye trace represents moving to the right. **(C)** Correct rate of sequence performance before and after electrophysiological recording. A successful sequence is defined by finishing a visually-guided sequence (four center-out saccades in the order of “LLRR”) and a memory-guided sequence. **(D)** The execution time of the whole sequence (“Seq duration” defined in **Figure 1B**) decreased during the sequence learning. **(E)** The coefficient of variation (CV) of execution time decreased with training. Early: training period before electrophysiological recording; Late: electrophysiological recording period. One session contained 20 – 50 trials.

##### Single Memory Saccade Task

Monkeys were also trained to perform a traditional delayed saccades guided by memory (Hikosaka O and RH Wurtz 1983; Bruce CJ and ME Goldberg 1985), as shown in **Figure 2-1B**. A fixation spot (0.3°×0.3°) first appeared at the center of the screen. After monkey maintaining fixation for 500 ms, a target spot (1.0°×1.0°) appeared at one of eight possible locations (7° relative to the screen center) throughout periphery of the visual field, then disappeared after 200 ms. A delay period began and lasted for 800-1,100 ms. At the end of the delay, the fixation spot disappeared, cueing the monkeys to initiate a saccade to the location of the vanished target within 500 ms. If the monkey reached the target area, the target (0.3°×0.3°) reappeared at the same location and the monkey was rewarded for maintaining fixation at this target for at least 300 ms.

#### Electrophysiological Recording

FEF neurons were extracellularly recorded by using single-unit tungsten microelectrodes (FHC; tip diameter 3 μm, impedance 1–2 MΩ). A hydraulic microdrive (FHC, USA) was implemented to advance the microelectrode into the cortex through a transdural guide tube. Neural signals were collected with four steps: amplified, bandpass filtered (400 - 5,000Hz), digitized and recorded (AlphaLab SnR, Israel). We isolated spike times with online sorting module (AlphaLab SnR, Israel) or offline sorting software Spike2 V8 (Cambridge Electronic) by using template matching algorithm.

FEF was identified using a combination of magnetic resonance imaging, white/gray matter transitions and evoked saccadic eye movements by microstimulation (**Figure 2**). Usually, saccadic eye movements were evoked from the anterior bank of arcuate sulcus. Penetrations where the typical ballistic eye movements were observed for at least 50% of microstimulation trials (200 Hz biphasic pulse trains, pulse width: 200 μs, interpulse interval: 100 μs, 50~150 ms duration, current magnitude: 30-100 μA; AlphaLab SnR) were defined as FEF.

**Figure 2.**
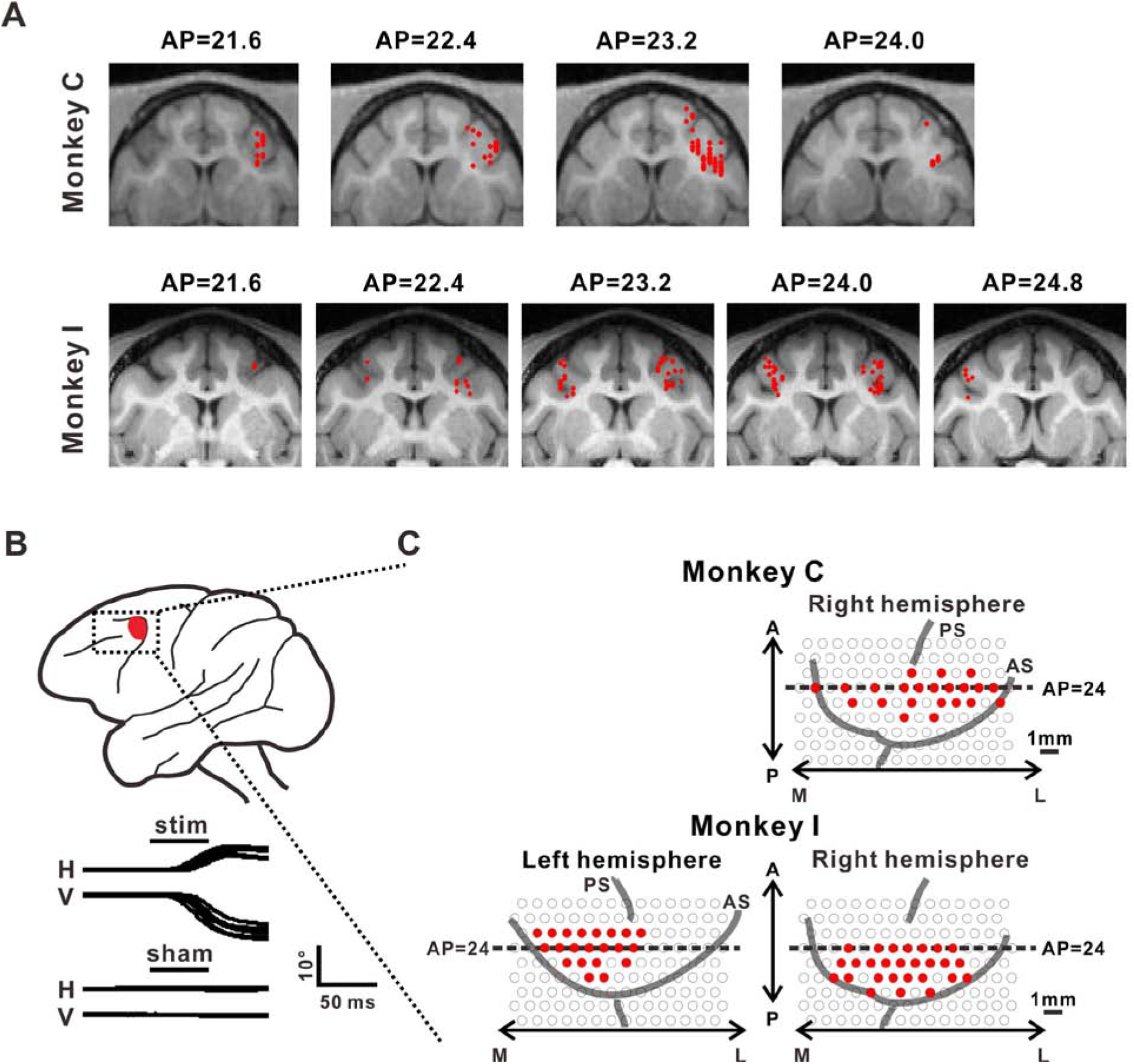
Anatomical localization of recording sites in monkey C and monkey I. **(A)** Neural recording sites was identified by magnetic resonance imaging. Recording sites of 2 monkeys were overlaid on magnetic resonance images (upper panels: Monkey C, n = 117 cells; lower panels: Monkey I, n = 109 cells). Coronal sections, spaced 0.8 mm apart, are shown from posterior to anterior; cells (marked with red symbols) located within 0.8 mm of each section were projected onto that section. **(B)** FEF area was also verified by electrical microstimulation. Examples of 50 μA and sham stimulus at FEF (marked with red symbols) were presented. **(C)** The penetration sites of FEF on the recording grid, marked with red filled circles.

#### Sub-threshold and supra-threshold Microstimulation

The electrical stimulation was delivered through a microelectrode placed in FEF where strong response and consistent saccade tuning were observed, and we used similar stimulus mode as above mapping procedure but with 200 ms duration and different amplitude (sub-threshold: 20 μA; supra-threshold: 100-200 μA) at the time when monkey settled on the fixation window and was about to perform the forthcoming saccade. Thus, in sub-threshold microstimulation experiment, monkeys were still able to finish the sequential saccades smoothly and the accuracy of performance did not be hampered. For sub-threshold microstimulation, one experiment block contained 250 trials, 80% of which were with stimulation and 20% were sham controls. Stimulations on each combination of 4 numerical orders and 2 contexts (visually-guided and memory-guided) were randomly interleaved so that animals could not predict which action within the sequence will be stimulated, resulting 25 repetitions for one saccade condition on average. Similar procedure was carried for supra-threshold microstimulation, but only 20% of trials with stimulation. Thus, one experiment block contained more than 400 trials to keep at least 10 repetitions.

#### Reversible inactivation

To further exmaine the causal role of FEF on sequential saccade, we performed the reversible chemical inactivation with muscimol (a GABA_A_ agonist), by using a “microinjectrode” (Chen LL et al. 2001; Chowdhury SA and GC DeAngelis 2008; Gu Y et al. 2012; Chen A *et al*. 2016). At the same time, we monitored multi-unit activities. We slowly inject 2 μl (10 μg/μl) of muscimol over ~20 minutes using a minipump, which allowing to inactivate a roughly spherical region with a radius of ~2 mm for several hours of duration (Arikan R et al. 2002). Performance on each task is measured both before and after the injection.

### QUANTIFICATION AND STATISTICAL ANALYSES

All analyses were performed using custom scripts written in MATLAB (Mathworks, Natick, MA). 226 sequence task related neurons were selected (80%, p < 0.05 in Wilcoxon rank-sum test compared with baseline fixation response). Among these cells, 165 cells were recorded in both sequence and single memory saccade tasks, which were analyzed for comparison of single and sequential saccades. Peri-event time histograms (PETHs) was constructed using 5-ms time bins. Neuronal responses of 200 ms after visual stimulus and ±100 ms from saccadic onset were used to calculate visual and saccadic responses, respectively. For data acquired from memory-guided saccade task, a peak response vector was yielded from PETH, which contains the maximal response across directions at each point in time. We performed Rayleigh test to assess the statistical significance of direction selectivity in visual and saccadic responses and the preferred direction was the direction of peak response vector summation.

For sequential saccade task, we defined three indexes to describe how different levels of sequence were encoded. Direction index (DI) was given by:

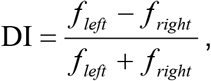

where f_left_ and f_right_ are the average responses from saccades towards left and right directions, respectively. Start index (SI) was given by:

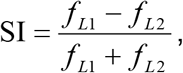

where f_L1_ and f_L2_ are the average responses from the first and second leftward saccades in the sequence, respectively. End index (EI) was given by:

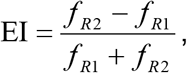

where f_R2_ and f_R1_ are the average responses from the last two rightward saccades, respectively. Memory index (MI) was given by:

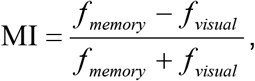

where f_memory_ and f_visual_ are the average responses from the memory- and visually-guided sequence, respectively.

Occasionally animals would perform a longer sequential saccades containing the required L-L-R-R order, such as L-L-L-R-R. In this case, neural activity and behavioral data for the first L, second L, second to final R and final R were analyzed. Results are presented as mean ± SEM unless otherwise stated, the significance level is 0.05.

### DATA AND SOFTWARE AVAILABILITY

Custom code for analyses will be provided upon reasonable request to the Lead Contact.

## Results

### Behavioral learning of saccade sequences in monkeys

To investigate whether and how FEF contributes to sequential saccades, we trained two rhesus monkeys (C and I) to perform four center-out saccades in the order of Left-Left-Right-Right (named “LLRR”) that were either visually-guided or memory-guided (see Materials and Methods for details; **Figure 1A**). **Figure 1B** shows the eye movement during a visually-guided sequence followed by a memory-guided sequence during electrophysiological recordings. It should be noted that the initiation and termination of each saccade was largely self-determined, thus there was no requirement for maintaining fixation during the multiple saccades (**Figure 1-1**). Monkeys were rewarded only if they correctly finished the sequence of four center-out saccades according to the LLRR order under visually-guided or memory-guided conditions.

Through training, the percentage of successful sequence (measured as “correct rate”) gradually increased across sessions (**Figure 1C**). When the correct rate of performance reached and maintained a level higher than 75%, we started electrophysiological recordings (marked with a dashed line in **Figure 1C**). At the same time, the duration of time required to perform the whole sequence of saccade (named “seq duration”, consisting of both visually- and memory-guided sequence) at the late stage (mean ± SD: 5355 ± 32 ms) was significantly shorter than the early stage (**Figure 1D**; mean ± SD: 6475 ± 44 ms, p < 0.01 in Wilcoxon signed rank test). Similarly, the coefficient of variation (CV) of sequence duration also decreased (**Figure 1E**; early: 0.067 ± 0.002, late: 0.046 ± 0.001, p < 0.01 in Wilcoxon signed rank test). These results indicate that the monkeys can learn the LLRR sequence, demonstrating quicker, more robust and stereotyped sequence performance with training.

### FEF neurons showed distinct direction preference during sequential vs. single saccades

Guided with the MRI scans (**Figure 2A**) and microstimulation (**Figure 2B**), we recorded from well isolated single neurons in FEF without prescreening, and collected activity from totally 226 (Monkey C: 117, Monkey I: 109) FEF neurons during the LLRR sequential saccade task. These neurons were recorded from the right hemisphere of monkey C (n = 117), left (n = 41) and right (n = 68) hemisphere of monkey I, respectively (**Figure 2A**). The relevant penetrations (marked with red dots) on the recording grids were shown in **Figure 2C**. It should be noted that for most recording sites (Monkey C: n = 111; Monkey I: n = 54), we also confirmed the physiological activities with a traditional single memory saccade task (**Figure 2-1A**) and classified these neurons into visual (6%, **Figure 2-1B**), visuomotor (20%, **Figure 2-1C**) and motor neurons (74%, **Figure 2-1D**) according to their visual and saccadic responses (**Figure 2-1E, F**).

Traditionally, FEF was considered to play a major and simple role in determining the direction of single forthcoming saccades (Bruce CJ and ME Goldberg 1985; Bruce CJ et al. 1985; Schall JD 1991; Isoda M and J Tanji 2003). Since most cells were tested with both single memory saccade and sequential saccade tasks, we first examined whether the saccade selectivity was maintained during the sequential task. **Figure 3A** shows an example FEF neuron with strong direction preference to the leftward saccade during single memory saccade task. Similar preference was observed when monkey was performing sequential saccade tasks (**Figure 3A**, right panel), by comparing the firing rate preceding saccades in the left and right directions. This type of neurons was classified as “consistent” cells. Interestingly, we also observed some cells exhibited different direction preference between these two tasks, as shown in **Figure 3B**. This cell had a leftward preference during single saccade task, but more activities were observed when monkey was making the 1^st^ rightward saccade during LLRR saccade sequence (right panel, **Figure 3B**). It should be noted that we verified the inconsistent direction preference by maintaining online good isolation from single neuron and also verified the spike waveform with offline analysis, as shown in the inserted right upper panels for the spike waveforms. Accordingly, such cell was classified as “inconsistent” cell.

**Figure 3.**
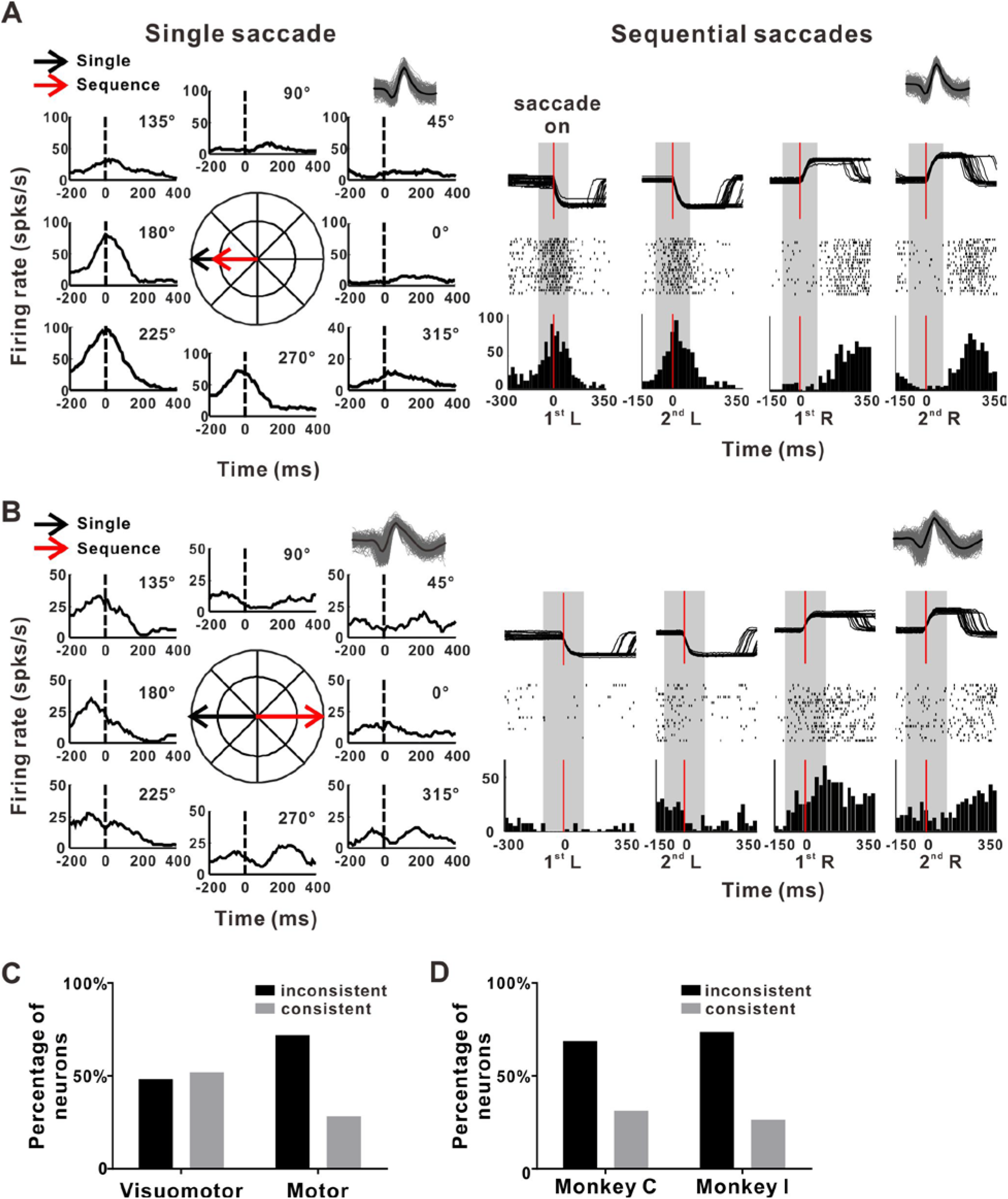
Dynamic encoding of saccade direction in FEF. **(A)** The FEF example neuron with consistent direction preference in the single memory-guided saccade task (left) and sequential saccade task (right). Same neuron recorded in two tasks was verified by the waveforms of spikes (insets). Saccade on markers are shown by vertical lines. **(B)** An example neuron with inconsistent direction preference during two different tasks. **(C)** The distribution of consistent and inconsistent cells in visuomotor and motor groups. n = 165 neurons, Wilcoxon signed rank test. **(D)** Similar proportions of consistency in two monkeys.

Among 155 cells recorded from FEF with significant leftward or rightward saccade preference, around two-thirds of neurons (102/155) exhibited “inconsistent” directional preference in sequence condition compared with the single saccade condition. The percentage of “inconsistent” cells was higher in motor cells (71.9%, n = 87 of 122 cells) than in visuomotor cells (48.1%, n = 15 of 33 cells, p = 0.0062 in Fisher’s exact test; **Figure 3C**). The proportion of “inconsistent” vs. “consistent” cells was similar in two monkeys and there was no individual difference (p = 0.79 in Fisher’s exact test **Figure 3D**). Since the saccadic direction and amplitude in single memory saccade were exactly the same as in sequential saccades (7 degree of visual angle; see Methods), the large proportion of “inconsistent” cells in FEF indicates that FEF might be involved in dynamic encoding of the forthcoming saccade direction depending on the previous saccade history.

### FEF neurons encode sequence structure

We then investigate whether FEF neurons encode the sequential-related information beyond the direction of single saccade. To address this question, FEF neuron activities during each saccade of the LLRR sequence were analyzed at different levels (element, subsequence and sequence) of sequential structure (*Geddes CE et al*. 2018). Consistent with previous study (Isoda M and J Tanji 2003), we did observe that some FEF neurons only respond to specific direction, as shown in **Figure 4A-C**. An example with stronger firing rate to the leftward saccades (61 spks/s) than to the rightward saccades (8 spks/s) (p = 1.2*10^−14^, paired t test) was shown in **Figure 4A**, and there is no significant difference between two left saccades (p = 0.93, one-way ANOVA with Tukey’s multiple comparisons test) or two right saccades (p = 0.082, one-way ANOVA with Tukey’s multiple comparisons test). As a comparison, **Figure 4B** shows another example FEF neuron with stronger firing rate to the rightward saccades. Among the population cells recorded from FEF, about half (45.9%, n = 95) neurons showed such activation related to each individual saccade direction within the sequence (here we called “Element” cell). Among these element cells, 36 (37.9%) neurons preferred to the left saccades, 23 (24.2%) neurons selectively tuned to the right saccades and the rest 36 (37.9%) neurons do not show significant preference to the leftward or rightward saccades (**Figure 4C**).

**Figure 4.**
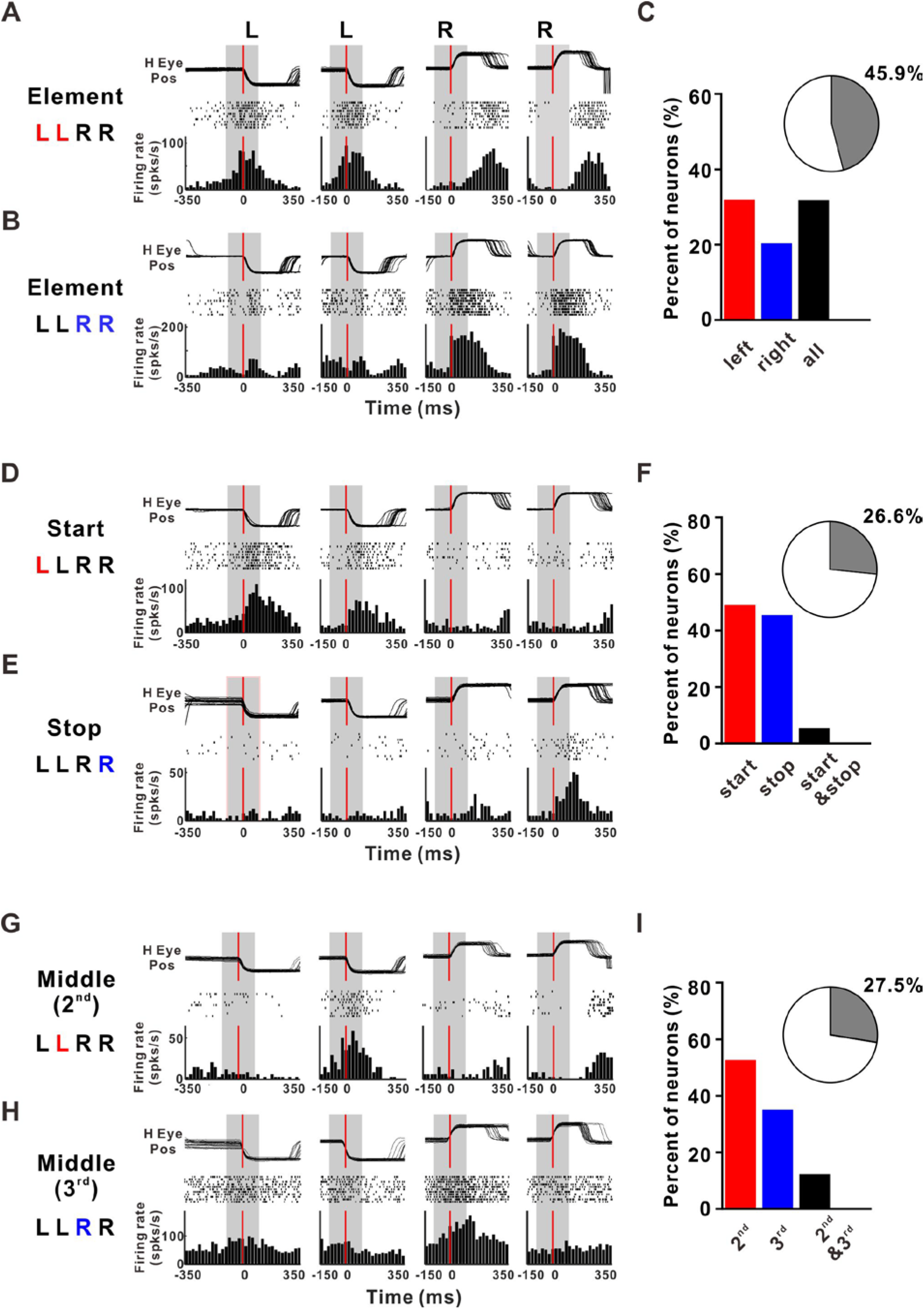
FEF neurons encode sequence structure. **(A-B)** Example neurons with responses to each preferred element. **(A)** show the activities of element example preferred leftward saccade during four saccades (Left, Left, Right, Right) within the sequence: Top: the superimposed eye traces from 20 repetitions, aligned to the initiation of the 1^st^ leftward saccade, 2^nd^ leftward saccade, 1^st^ rightward saccade, and 2^nd^ rightward saccade; Middle: each dash indicates a spike. Bottom: Peri-event time histogram (PETH) of the same neuron. **(B)** Another element example with preference to the rightward saccade. **(C)** Proportion of FEF showing element activity. The percentage among the whole recorded FEF was presented as pie plot in the inset. **(D-E)** Example neurons with start **(D)** and stop activity **(E)**. **(F)** Proportion of FEF showing sequence start/stop activity. **(G-H)** Example neurons with subsequence switch-related activity (**G**: example preferred to the end of left-subsequence; **H**: example preferred to the start of right-subsequence). **(I)** Proportion of FEF showing subsequence switch-related activity.

However, nearly a quarter (26.6%, n = 55) FEF cells showed sequence-level start/stop-related activity, as shown in **Figure 4D-F**. A start example was shown in **Figure 4D**, with clear preference to the first left saccade. To quantify the involvement of FEF in higher structural level of a learned sequence, we defined a Start Index (SI) to indicate the rank selectivity by comparing the difference of neural responses across these two left saccades ((f_L1_−f_L2_)/(f_L1_+f_L2_), see Methods for details). For this example, the SI was 0.25 (p = 2.3*10^−4^, one-way ANOVA with Tukey’s multiple comparisons test). **Figure 4E** illustrated an example of stop cell, with higher firing rates to the last rightward saccade than to the first rightward saccade. Similarly, we also defined an End Index (EI) by comparing the responses to the two right saccades ((f_R2_−f_R1_)/(f_R2_+f_R1_), see Methods), and EI for this example is 0.46 (p = 0.0019, one-way ANOVA with Tukey’s multiple comparisons test). Across the population, the proportions of FEF neurons preferred “start” and “stop” were close (49.1% vs. 45.5%, **Figure 4F**). Besides, we also observed a few cells (n = 3, 5.4%) with both preference to the first leftward saccade and the last rightward saccade, named “start & stop” cell.

Another quarter (n = 57, 27.5%) FEF neurons were selectively active during the transition from the left to the right subsequences (**Figure 4G-I**), with stronger firing rate to the 2^nd^ saccade (or the last saccade for the left subsequence, see the example in the upper row in **Figure 4G**) or the 3^rd^ saccade in the sequence (or the first saccade for the right subsequence, see the example in the bottom row in **Figure 4H**). Among the population, the percentage of cells with preference to the 2^nd^ saccade (52.6%) is slightly higher than that preferred to 3^rd^ saccade (35.1%). A few cells (n = 7, about 12%) with preference to both 2^nd^ and 3^rd^ saccade in the sequence (**Figure 4I**).

Together, these data suggest that FEF not only encode specific saccade direction but also exhibit strong sensitivity to the structure of sequence. In other words, FEF can be dynamically involved in encoding both the direction and the abstract structure of saccade sequences at element, subsequence and sequence of all three levels. It should be noted that when we further seperate the analysis window into pre saccade (−100 ms ~ 0, **Figure 4-1A-C**) or post saccade (0 ~100 ms, **Figure 4-1D-F**), similar percentages of rank activities were still observed.

### Context-dependent sequence encoding in FEF

To further demonstrate the dynamic coding in FEF, we also compared the activities of the same FEF neurons between visually- and memory-guided sequential saccades, since saccade movements can be guided by external and internal memory and both factors are important for sequence learning and execution under different environmental conditions (MacAvoy MG and CJ Bruce 1989; Pierrot-Deseilligny C 1991). **Figure 5A** illustrates the firing activity of an example FEF neuron during the performance of visually-guided sequence and memory-guided sequence. It is evident that the neuron exhibited strong firing activity during the switch from the LL to the RR subsequence during memory-but not visually-guided sequential saccades, although this cell had similar rank activities for both tasks (preferring to 3^rd^ saccade or 1^st^ rightward saccade in the current sequence). We also found that some FEF neurons changed the rank activities, as illustrated in **Figure 5B** (for a start cell in memory sequence but not in visually task) and **Figure 5C** (for an end cell during visually sequence but with middle activities during memory task). To quantify the dynamic encoding characteristics reflected by difference between these two tasks, we computed the memory index (MI) ((f_memory_−f_visual_)/(f_memory_+f_visual_), for details see Methods) in which MI = 1 means the neuron only responds during memory-but not visually-guided sequence and MI = −1 means vice versa. For this particular cell in **Figure 5A**), the MI is 0.47 (p = 2.5*10^−8^, paired t test). Although there was no significant difference between the two modes of sequence at the behavior level (p > 0.05 in Wilcoxon rank-sum test; **Figure 1-1**), 69% (156/226) of FEF neurons demonstrated context-dependent characteristics (**Figures 5D**), with significantly (p < 0.05, paired t test) different firing activities for the same action element in the sequence between visually-guided and memory-guided task. Among these cells, 35.63% showed difference for the 1^st^ L saccade between visually- and memory-guided sequential saccade tasks, while 26.12%, 18.54% and 19.71% fired differently between these two contexts for the rest 2^nd^ L, 1^st^ R and 2^nd^ R saccade respectively. It should be noted that the percentages of overlapped action positions were divided equally in the percentage calculation (**Figure 5E**).

**Figure 5.**
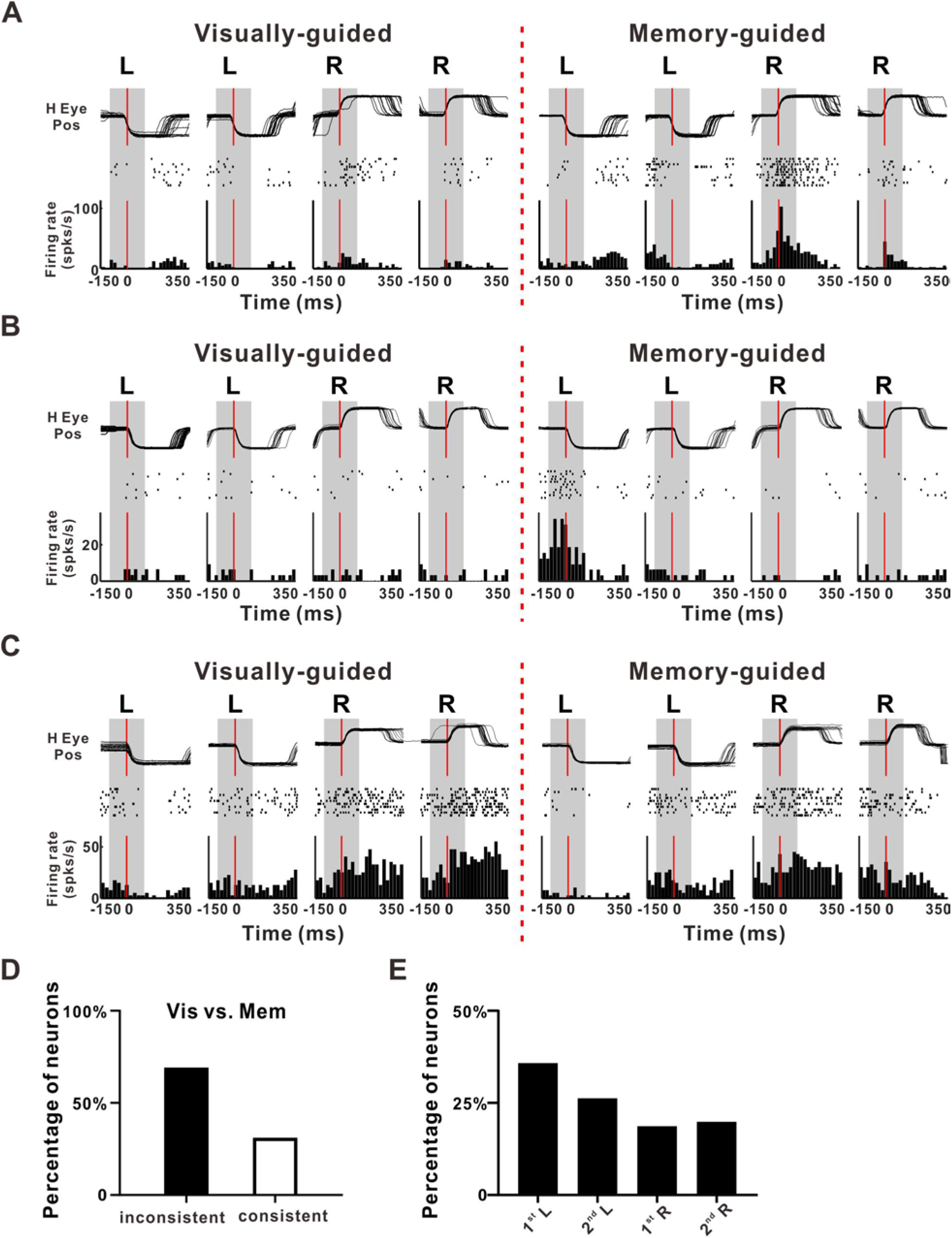
Context-dependent sequence-related neuronal activity in FEF. **(A)** A representative neuron exhibiting different activity in visually- and memory-guided contexts. **(B)** An example neuron with start activity in memory-guided sequence but not in visually-guided sequence. **(C)** An example neuron with clear end activity in visually-guided sequence but with middle activities (stronger responses to 3^rd^ saccade in the sequence) during memory-guided task. **(D)** The percentage of cells with (black bar, n = 156 cells) or without (gray bar, n = 70 cells) significantly different firing during visually- vs. memory-guided sequence saccade task (p < 0.05, paired t test). **(E)** The percentage of cells with significantly different firing between two contexts at different action positions within sequence (p < 0.05, paired t test).

Thus, FEF neurons showed clear difference between these two types of sequence tasks, suggesting that FEF controls saccade movements by encoding the combination of sequence structure and specific context in a manner much more complicated than previously thought.

### Effects of FEF microstimulation on learned sequence behavior

To test the causal link between FEF activity and sequential saccades, we applied supra-threshold microstimulation (for details, see Materials and Method) when monkey was performing sequential task to some sites in left and right FEF marked with red dots in **Figure 6A** and **Figure 6B** respectively. For these sites, we can easily use microstimulation to evoke saccade response when monkey was maintaining fixation only. However, it is interesting to note that when we applied same microstimulation as we did in fixation task, the percentage of successfully induced saccade was reduced when monkey was performing memory-guided sequence for both left (**Figure 6C**) and right hemisphere (**Figure 6D**). For 87% trials, saccade can be induced successfully before the 1^st^ action of the sequence, and this percentage is lower during the middle of sequence, with 78%, 38% and 86% for the time point before 2^nd^, 3^rd^ and 4^th^ saccade of the sequence (**Figure 6C**). Similar results were observed in the right FEF, as shown in **Figure 6D**, indicating that the threshold to generate a saccade with microstimulation is different for the various positions during the sequence performance.

**Figure 6.**
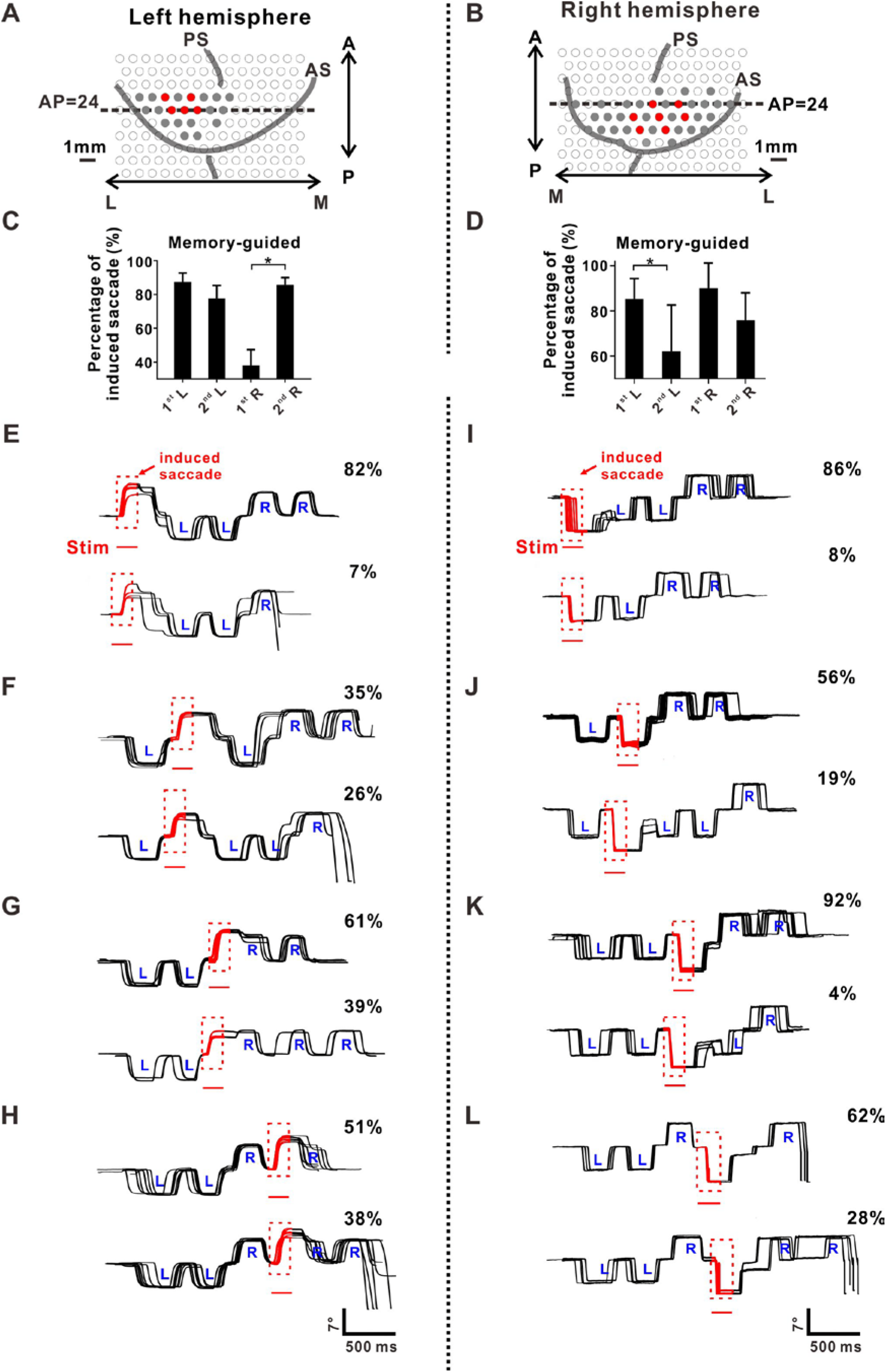
Effects of supra-threshold microstimulation of FEF on sequential saccades. **(A-B)** Location of all the penetrations for supra-threshold microstimulation in left **(A)** and right **(B)** hemisphere on the recording grid, marked with red filled circles. **(C)** The percentages of successfully evoked saccade by stimulation at different time point within memory-guided sequence for left hemisphere (n = 14 penetrations, 1^st^ L vs. 2^nd^ L, p = 0.21; 1^st^ R vs. 2^nd^ R, p = 2.4*10^−5^, paired t test). **(D)** The percentages of successfully evoked saccade for right hemisphere (n = 6 penetrations, 1^st^ L vs. 2^nd^ L, p = 0.028; 1^st^ R vs. 2^nd^ R, p = 0.21, paired t test). Formats same as left hemisphere **(C)**. **(E-H)** Example eye traces obtained during supra-threshold stimulation of left FEF sites at different time point (**E**: stimulation before the first saccade (1^st^ L); **F**: stimulation before the 2^nd^ saccade (2^nd^ L); **G**: stimulation before the 3^rd^ saccade (or 1^st^ R); **H**: stimulation before the 4^th^ saccade or 2^nd^ R) within sequence. Red traces represented the eye traces evoked by electrical stimulation. **(I-L)** The effect of stimulation from the right hemisphere. Formats are same as the left hemisphere. For all figures, *p < 0.05, paired t test. Error bars denote SEM. See also Figure 4-1.

**Figure 6E-H** and **6I-L** show how the supra-threshold microstimulation affects the sequence behavior in left and right FEF respectively. Specifically, for rightward saccade induced by the supra-threshold microstimulation before the first saccade of the sequence, the monkey properly initiated and finished the whole sequence of LLRR without any difficulty (**Figure 6E**). Such type of behavior occurred in most (82%) of the trials with simulation successfully evoked saccade. For the second most (7%) trials, after the induced saccade by microstimulation, monkeys finished the two left saccades and one right saccade, then broke the sequence (for the rest trials, see **Figure 6-1A**). When the rightward saccade was induced before the 2^nd^ left saccade of the sequence, the monkey ignored the evoked saccade and continued to finish the correct rest of the sequence for about 35% trials (upper panel, **Figure 6F**), and reset the sequence for about 26% trials (lower panel, **Figure 6F**) (for the rest trials, see **Figure 6-1B**). These data suggested that the monkey could distinguish the saccade elicited by the external FEF stimulation from those internally generated, and confirmed that the learned action sequences are not organized in a serial chain but likley in a hierarchy (*Geddes CE et al*. 2018). When the rightward saccade was evoked before the 1^st^ or 2^nd^ right saccade in the sequence, the monkey usually adjusted the eye position to the proper position after the induced saccade, so that the saccade was successfully integrated to the overall sequence (**Figure 6G** & **6H**, respectively). **Figure 6I-L** showed the results of sequence action after the left saccade was induced from the right hemisphere microstimulation at different time point. Overall, the induced saccade can be either ignored or adjusted to be integrated into the overall sequence without significant disruption of the learned sequence structure. Beside this supra-threshold microstimulation, we also tried sub-threshold microstimulation in FEF, but no clear significant effect was observed on the execution of sequential saccades (see **Figure 6-2 and 6-3**): monkey did not break or stop the sequential saccade task during stimulation, and only saccade latency increased for a few sites (**Figure 6-2D, and 6-3A-B**). These findings together suggest that perturbing FEF activity in any sequence position could evoke saccade but does not affect the following sequence execution, implying a hierarchical behavioral organization of learned saccade sequences (Geddes CE *et al*. 2018).

### Effects of inactivation of FEF on sequential saccades

Since FEF is part of a network of cortical and subcortical areas that controls eye movements, microstimulation only in FEF without change the structure of sequential saccade does not tell the exact role of FEF during sequential saccade. To further explore a clear function of FEF in sequential saccades, we turned to reversible chemical inactivation experiments to analysis the deficit in oculomotor performance. Thus, we placed the injections in different sites of the FEF representing different saccade vectors. Two example inactivation sites were illustrated in **Figure 7A**: stimulation of the site in the left hemisphere before injection consistently elicited an up-rightward saccade indicated by the red arrow; stimulation of the right hemisphere site evoked down-leftward saccade indicated by the blue arrow. Neural activity surrounding the injection cannula tip was typically silenced by the end of the 20 min injection period, confirming the inactivation effect after successful delivery of the drug (right panel, **Figure 7A**).

**Figure 7.**
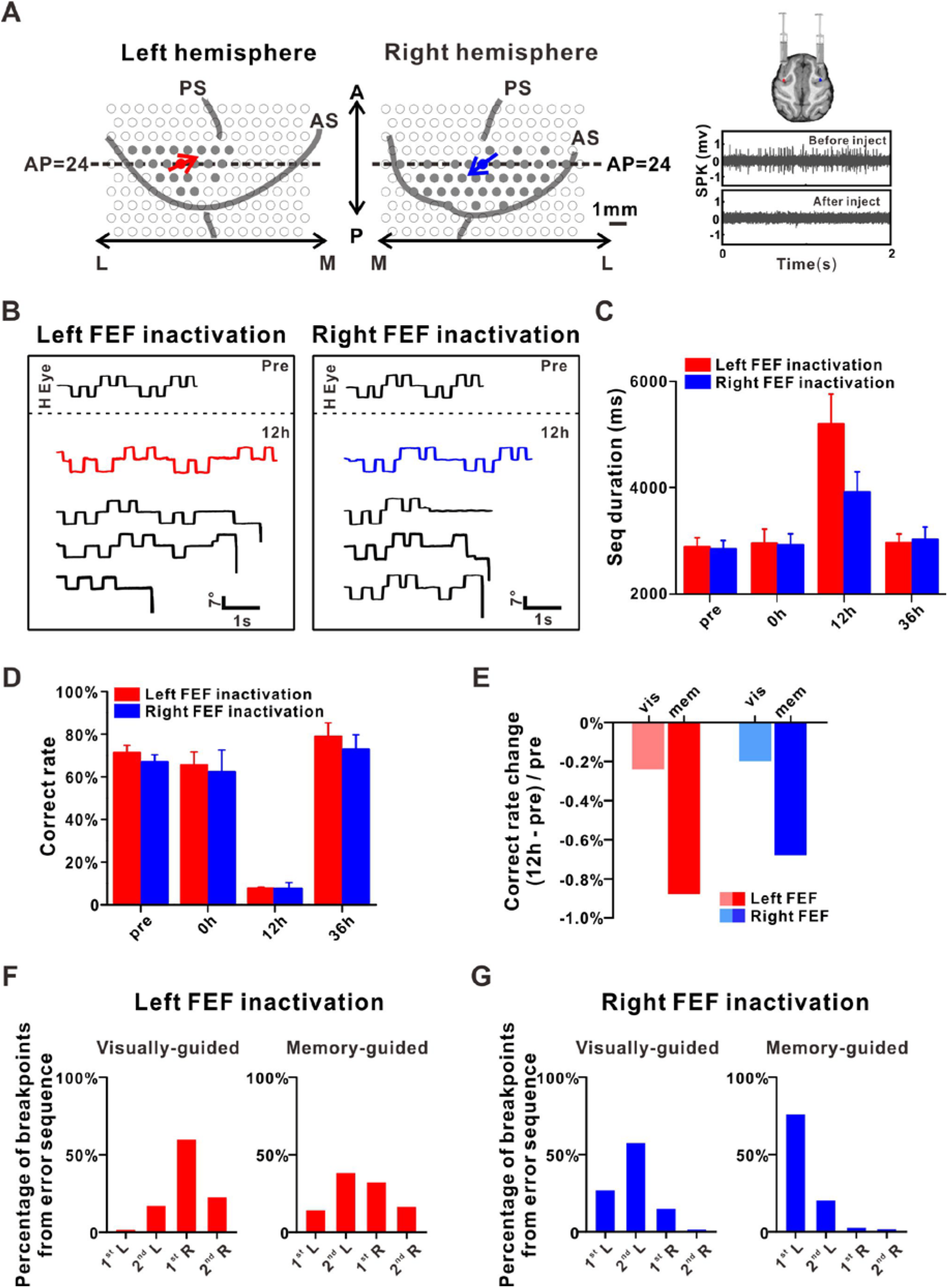
Effects of muscimol inactivation of FEF on overall task performance. **(A)** Location of the two example injection sites. Red and blue arrows indicate the saccade direction evoked by electrical stimulation at these two injection sites in left and right FEF respectively. The inset figures show that illustrations of muscimol injection in left and right FEF (upper), and the muscimol injection suppresses neural activity in a large region (lower). **(B)** Examples of sequential saccade performance at 12h time point after the muscimol injection compared with the control data (”Pre”): red (left FEF) and blue (right FEF) traces indicated the whole sequence was finished with an increased duration, while the black traces show some examples with sequence broken at different locations. **(C)** The effect on sequence duration at different time point after muscimol injection (Left hemisphere (Red bar): 12h versus pre/0h/36h, p = 1.1*10^−11^, p = 0.016 and p = 8.9*10^−5^, respectively; Right hemisphere (Blue bar): 12h versus pre/0h/36h, p = 2.2*10^−8^, p = 0.025, and p = 0.0056, respectively, two tailed t test). Only trials with whole sequence (4 center-out saccades in visually-guided together with 4 saccades in memory-guided task) finished were considered. **(D)** The correct rate of sequential performance after muscimol injection. The abscissa marks the four different time points at which data were collected: Pre, 0, 12, or 36 h (Red bars represent left FEF inactivation: 12h vs. pre/0h/36h, p = 2.4*10^−5^, p = 0.021 and p = 0.0028, respectively; Blue bars represent right FEF inactivation: 12h vs. pre/0h/36h, p = 3.2*10^−5^, p = 0.069,and p = 0.0055, respectively, two tailed t test). **(E)** The normalized correct rate change between after 12h and before (“pre”) inactivation, separately for visually-guided and memory-guided sequence task. **(F-G)** The percentage of trials broke at different sequence location after inactivation in left **(F)** and right **(G)** FEF. Muscimol data were analyzed using two tailed t test. Error bars denote SEM **(C)** and STD **(D)**, respectively.

For each inactivation experiment, control data were collected on the first two days before inactivation (“pre”). Immediately after the muscimol injections, a block of behavioral data was obtained (“0h”). Further two blocks of behavior trials were performed around 12h and 36h after inactivation, respectively. Results of several example eye traces after inactivation were compared with “pre” control in **Figure 7B**. Overall, monkey’s performance was severally affected around “12h”: for some trials, monkey can finish the whole sequence (red and blue traces for left and right hemisphere respectively), but the sequence duration was greatly increased (pre vs. 12h: 3835 ms vs. 5965 ms for left hemisphere; 3745 ms vs. 5115 ms for right hemisphere). For the majority of trials at 12h, monkey just break sequence, as shown for the black traces at 12h in **Figure 7B**.

For the trials with whole sequence finished, **Figure 7C** summarized the effect on sequence duration from 4 inactivation experiments. It is obvious to note that these trials was performed more slowly at 12h after inactivation, with averaged duration (Mean ± SD: left, 5200 ± 560 ms; right, 3912 ± 382 ms) significantly larger than “pre” control (Mean ± SD: left: 2883 ± 170 ms; right: 2839 ± 162 ms) (p = 1.1*10^−11^, two tailed t test). However, in most cases, saccade sequences were broken at 12h after muscimol injection (**Figure 7D**) with only 7.8 and 7.7% of correct rate for left and right hemisphere respectively, which are significantly lower than “pre” control (p = 2.4*10^−5^, two tailed t test). And the effect on the memory-guided sequence was more severely than the visually-guided sequence, as shown in **Figure 7E**. For those error trials (monkey did not finish the whole sequence), we also checked how the inactivation affects the actions at different sequence location, as shown in **Figure 7F**. For visually-guided task, most sequence was broken at 1^st^ rightward saccade (1^st^ R) after left FEF inactivation, while most sequence stopped at 2^nd^ leftward saccade (2^nd^ L) during memory-guided task. In addition, the same action at different rank position within sequence was affected differently. As shown in **Figure 7F**, it is obvious that the 1^st^ leftward saccade (vis: 1.4%; mem: 14.1%) was less affected than the 2^nd^ leftward saccade (vis: 16.9%; mem: 38.0%) in the left hemisphere. On the contrary, the 1^st^ rightward saccade (vis: 59.5%; mem: 31.8%) was affected more than the 2^nd^ rightward saccade (vis: 22.3%; mem: 16.1%). For the right hemisphere, we can also see that same actions at different sequence location was affected differently for both visually-guided and memory-guided context with 26.7% for 1^st^ and 57.3% for 2^nd^ leftward saccade during visually-guided sequence, and with 75.8% for 1^st^ and 20.2% for 2^nd^ leftward saccade during memory-guided sequence, respectively (**Figure 7G**). These results strongly indicate that FEF inactivation impaired sequence not solely due to disrupting single saccades.

Although FEF inactivation with muscimol usually affects both visually- and memory-guided saccades, **Figure 7E** shows the effect to the memory-guided saccades was different from visually-guided sequence. To make a quantified comparison between these two different sequence tasks, we further checked the saccade latency, and found the inactivation effect on the memory-guided saccades was more severe for both single saccade task (for details, see **Figure 8-1A**) and sequential saccade task (**Figure 8**). Since the ipsilateral saccades were not as strongly affected by the injection as the contralateral saccades, here we only show the example of rightward saccade for left hemisphere in **Figure 8A**. It is notable that the saccade latencies were clearly increased for all the rightward saccades after injection. The population results were summarized in **Figure 8B**. It is interesting to note that the inactivation affects the sequential saccades task more than the single saccade task, with averaged increased latency (127.2 ms for visual and 198.3 ms for memory) in sequential saccade task significantly larger than 46.4 ms in single saccade task (p = 0.0062, two tailed t test). In addition, even for the same saccade action during sequential saccades task, the inactivation effect was different between the two leftward or rightward saccades. As shown in **Figure 8B**, the increased saccade latencies for the 2^nd^ rightward saccade averaged 179.0 ms, larger than the 1^st^ rightward saccade (132.6 ms) during visually-guided sequence, although not significant (p = 0.18, one-way ANOVA with Tukey’s multiple comparisons test). Opposite effects were observed in the memory-guided sequence, as shown in the right panel of **Figure 8B**, with mean value for the 1^st^ rightward saccade (325.3 ms) significantly larger than the 2^nd^ rightward saccade (153.0 ms) (p = 0.013, one-way ANOVA with Tukey’s multiple comparisons test). These again strongly suggest that FEF dynamically encodes the sequential saccades and the same saccades in the different position of a sequence are differently affected by FEF inactivation. **Figure 8C** summarized the inactivation effects from population at different time point. Again, the strongest effect was observed at 12h after inactivation for saccade latencies. Similar results were observed for the inactivation in the right hemisphere (**Figure 8D-F**), with stronger inactivation effects for the latencies of the left saccades during the memory-guided sequence and more obvious effects at 12h after inactivation. These findings were further confirmed in a control task where monkeys are trained to perform RRLL sequence, and inactivation has the qualitatively similar effect to the sequence (**Figure 8-1**).

**Figure 8.**
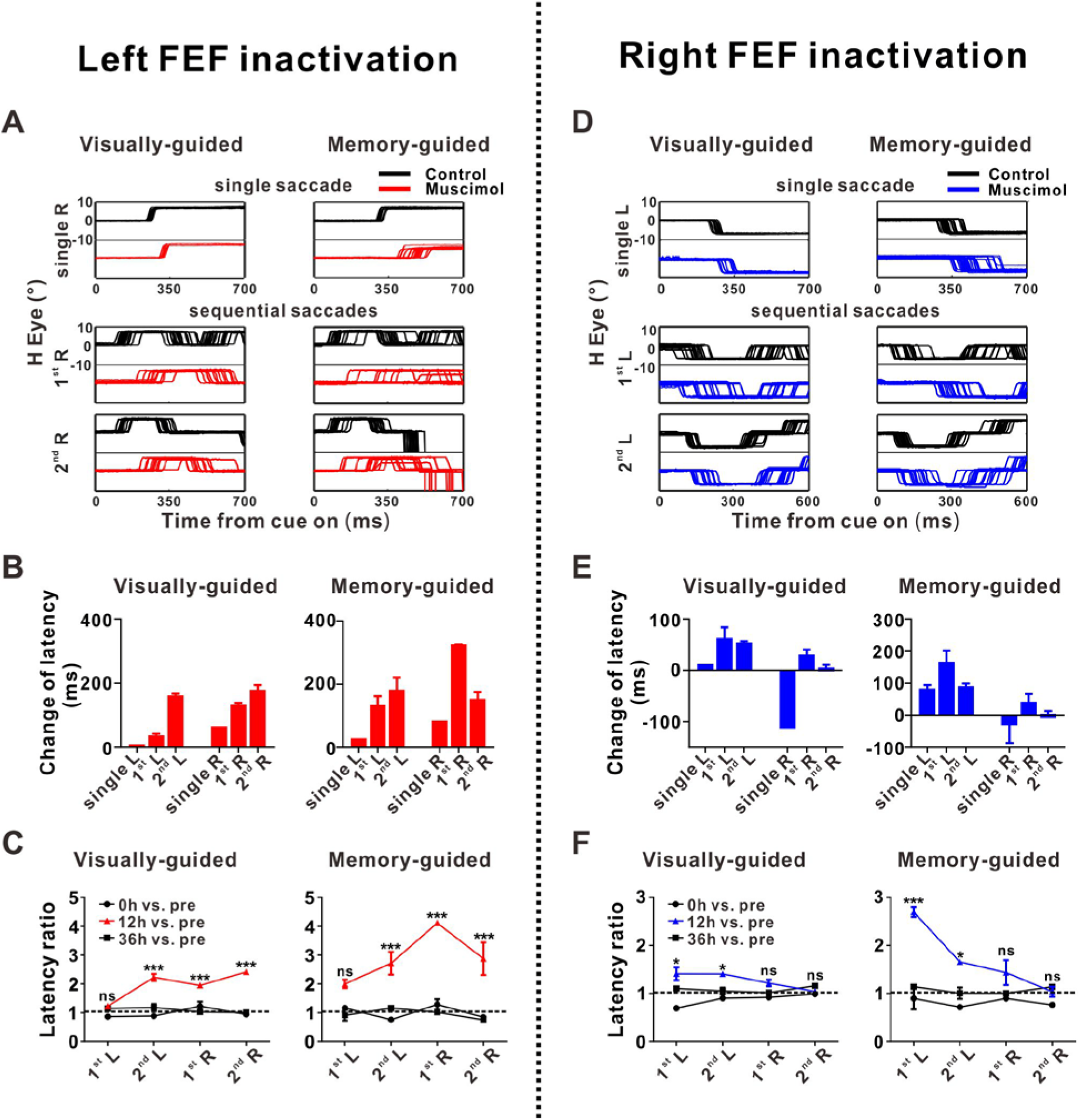
Effects of muscimol inactivation of FEF on sequential saccade performance. **(A)** Eye traces obtained during single saccade task and sequential saccade task at 12h time point after muscimol injections in left FEF (red traces) and control (black traces). **(B)** Summary of latency changes at 12h time point after injection. Visually-guided: 1^st^ L vs. 2^nd^ L, p = 0.23; 1^st^ R vs. 2^nd^ R, p = 0.18; Memory-guided: 1^st^ L vs. 2^nd^ L, p = 0.23; 1^st^ R vs. 2^nd^ R, p = 0.013, one-way ANOVA with Tukey’s multiple comparisons test. **(C)** Saccade latency ratio (saccade latency after injection divided by the latency measured before injection) changes as time. 12h vs. pre during visually-guided indicated by red line: 1^st^ L versus 2^nd^ L, p = 2.0*10^−4^; 1^st^ R versus 2^nd^ R, p = 0.084; 12h vs. pre (red line) during memory-guided: 1^st^ L versus 2^nd^ L, p = 0.34; 1^st^ R versus 2^nd^ R, p = 0.038, two-way ANOVA with Tukey’s multiple comparisons test. **(D-F)** Results from the muscimol injection in the right FEF. **(D)** Eye traces obtained during single saccade task and sequential saccade task at 12h time point after muscimol injections in right FEF (blue traces) and control (black traces). **(E)** Latency changes at 12h after injection in right FEF. Formats same as left hemisphere **(B)**. Visually-guided: 1^st^ L vs. 2^nd^ L, p = 0.23; 1^st^ R vs. 2^nd^ R, p = 0.31; Memory-guided: 1^st^ L vs. 2^nd^ L, p = 0.035; 1^st^ R vs. 2^nd^ R, p = 0.21, one-way ANOVA with Tukey’s multiple comparisons test. **(F)** Saccade latency ratio changes as time for right FEF inactivation. Formats same as **(C)**. 12h vs. pre during visually-guided indicated by blue symbols and line: 1^st^ L vs. 2^nd^ L, p > 0.9999; 1^st^ R vs. 2^nd^ R, p = 0.66; 12h vs. pre (blue line) during memory-guided: 1^st^ L vs. 2^nd^ L, p = 0.0028; 1^st^ R vs. 2^nd^ R, p = 0.38, two-way ANOVA with Tukey’s multiple comparisons test. For **(C)** and **(F)**, *p < 0.05, **p < 0.001, ***p < 0.0001, and ns, no significant difference with two-way ANOVA, followed by Sidak’s multiple comparisons test. Error bars denote SEM.

Together, these results suggest that inactivation of FEF affects the saccade differently depending on its position within the sequence, and the FEF activity may thus play a critical role in both organizing and execution of saccade sequences.

## Discussion

The FEF has traditionally been identified as an important area in saccade generation with low electrical currents (Bruce CJ and ME Goldberg 1985), thus the classic view for the role of FEF is simply encoding a saccade vector. However, by examining the activity of the same neuron under both single memory saccade and sequential saccades, we demonstrated that for more than 50% of FEF visuomotor and motor neurons (**Figure 3**), the direction selectivity during sequential saccades could not be simply inferred from single saccade behavior. This result suggests that the information encoding in FEF is more dynamic and complicated than previously thought.

Further analysis for sequential saccades turned out that although we observed high ratio of direction selective neurons in FEF (**Figure 4A** and **Figure 2-1**) which is consistent with previous study (Isoda M and J Tanji 2003), still a small part of FEF neurons exhibited sensitivity to the specific position/step of a saccade within sequence (**Figure 4D-I**). And the sequence-specific activity was more common when we only compared the same two left saccades (**Figure 4D, 4G**) or two right saccades (**Figure 4E, 4H**), indicating the same saccadic actions at different steps within sequence are not encoded in the same way by FEF. This was further confirmed by the inactivation results, with different error rate (**Figure 7F-G**) and different latency increased (**Figure 8B, 8E**, and **Figure 8-1D, F**) for the same action but at different steps within sequence. To some extent, these results strongly support some psychophysics theory about sequential saccades processed are not serial (Lu X *et al*. 2002; Isoda M and J Tanji 2003; McPeek RM et al. 2003; Schall JD 2004; Basu D and A Murthy 2020). Since most of analyzed FEF neurons were motor cells with significant perisaccadic activity, these dissimilar representations could not be simply explained by visual attention mechanisms. Instead, our results demonstrated that FEF encoded successive target location with embedding structural information such as start/stop state representations in sequential saccades, which could be useful in processing hierarchical actions (Hikosaka O et al. 1999; Jin X and RM Costa 2010; Jin X *et al*. 2014; Jin X and RM Costa 2015).

One interesting finding in our data is that the proportion of start or stop cells was greater than that of the sequential-related cells documented in a similar study in FEF (Isoda M and J Tanji 2003). The prominent differences may due to the complexity and temporal requirements of behavioral task. The action sequence used in this study is a self-paced task with symmetrical multi-layer structured, a commendable imitation of common motor activities. In addition, the sequence action was well consolidated with long-term training (see **Figure 1C-E**), so monkeys were more inclined to form a hierarchical structure of movement execution. This also agrees with previous studies in rodents (Jin X *et al*. 2014; Geddes CE *et al*. 2018), which proposed that the learned action sequences are organized in a hierarchy, and different hierarchical levels may be controlled by different brain regions or networks. Indeed, the proportion of sequence-level start/stop activity observed in FEF is lower than those previously observed in the striatum of mice trained with the similar action sequence (Geddes CE *et al*. 2018), suggesting the motor cortex (e.g., FEF in the current study) and basal ganglia might be operating at different levels of this hierarchy (Hikosaka O *et al*. 1999; Jin X and RM Costa 2010; Jin X *et al*. 2014; Jin X and RM Costa 2015). In fact, although FEF exhibited sequence-related neuronal activities (**Figure 4F, and 4I**), the additional saccade induced by microstimulation (**Figure 6**) was ignored by the monkeys during the execution of sequences. This result suggests that FEF appears to be either partly redundant, or complementary to, or acting in conjunction with other areas during the planning of sequential saccades, although it is clearly part of the network controlling the execution of saccades. And there are additional higher levels of hierarchical control for the sequence execution (Jin X and RM Costa 2015; Geddes CE *et al*. 2018), while the FEF likely acts as a lower level of controller for both planning and execution of sequence saccades. This is further supported by our muscimol experiment results, in which FEF inactivation dramatically disrupts the sequence execution at various positions within the sequence, beyond the impairment of single individual saccades (**Figure 7**). As a comparison, neurons in the striatum have been reported to show sequence-related neuronal activity during the execution of a learned LLRR sequence in mice (Geddes CE *et al*. 2018), and optogenetic stimulation of D1-expressing projection neurons in the striatal direct pathway facilitates the ongoing action within the sequence without altering the following sequence execution, like what are seen in the current study (Geddes CE *et al*. 2018). However, one noticeable difference is that the action induced by striatal direct pathway stimulation was taken into account by the animals to adjust the rest of sequence being performed (Geddes CE *et al*. 2018), whereas the saccade action evoked by FEF stimulation was largely ignored by the animals (**Figure 6**). Furthermore, ablation of striatal D1-expressing projection neurons significantly impairs overall sequence initiation (Geddes CE *et al*. 2018), in contrast with the effects of FEF inactivation specifically on subsequence (**Figures 7 and 8**). While we acknowledge that the two studies were performed in different species with mice vs. monkeys, it nevertheless implied again the possible functional difference between motor cortex and basal ganglia for operating at the different levels of hierarchical control for sequence execution (Jin X and RM Costa 2015; Geddes CE *et al*. 2018).

Nevertheless, our current study underscores the dynamic coding of saccadic parameters by FEF neurons, and reveals the underappreciated complexity and importance of FEF in controlling sequential saccades. Future studies should be aimed toward identifying the specific roles of frontal and higher-order motor cortices (Hikosaka O *et al*. 1999; Tanji J 2001; Lu X *et al*. 2002; Fujii N and AM Graybiel 2003; Shima K et al. 2007) as well as subcortical regions like thalamus and superior colliculus in the hierarchical control of learned saccade sequences.

## Acknowledgements

This work was supported by grants from National Basic Research Program of China (31871079) and Shanghai education committee of scientific research innovation (18JC1412500, 2018SHZDZX05) to A.C, and Shanghai Sailing Program (19YF1413500) to Z.P. We thank Drs. Tony Movshon, Jeff Erlich and SzeChai Kwok for helpful suggestions. We also feel grateful to Minhu Chen for computer programming.

## Author contributions

X.J. and A.C. designed the experiments. J.J., Z.P. and Q.W. conducted the experiments. J.J., Z.P., X.J. and A.C. analyzed the data and discussed the results. J.J. and Z.P. prepared the figures. J.J, Z.P., X.J. and A.C. wrote the paper.

## Extended Data

**Figure 1-1.**
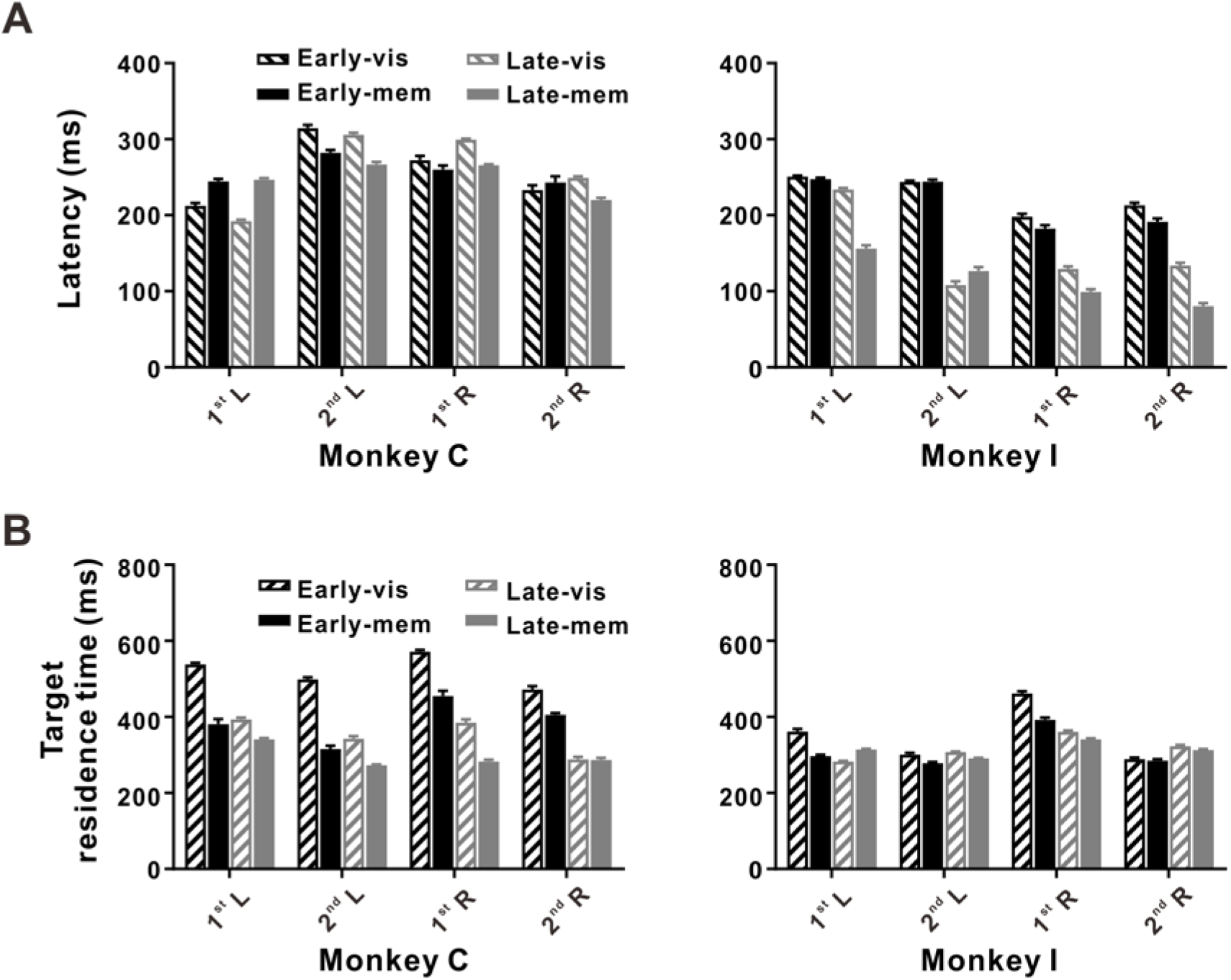
Summary of saccade latency and target residence time of individual saccade within sequential task. **(A)** Left panel: the summarized saccade latency from monkey C. Black and gray color indicate the early and late stage for performing sequential saccades respectively. Striped and filled bars represent the visually and memory-guided sequential task respectively. Right panel: The summarized results of saccade latency from monkey I. Formats same as left panel. **(B)** Left panel: the summarized target residence time (the time when monkey made a fixation at the target) from monkey C. right panel: results from monkey I. Error bars denote STD.

**Figure 2-1.**
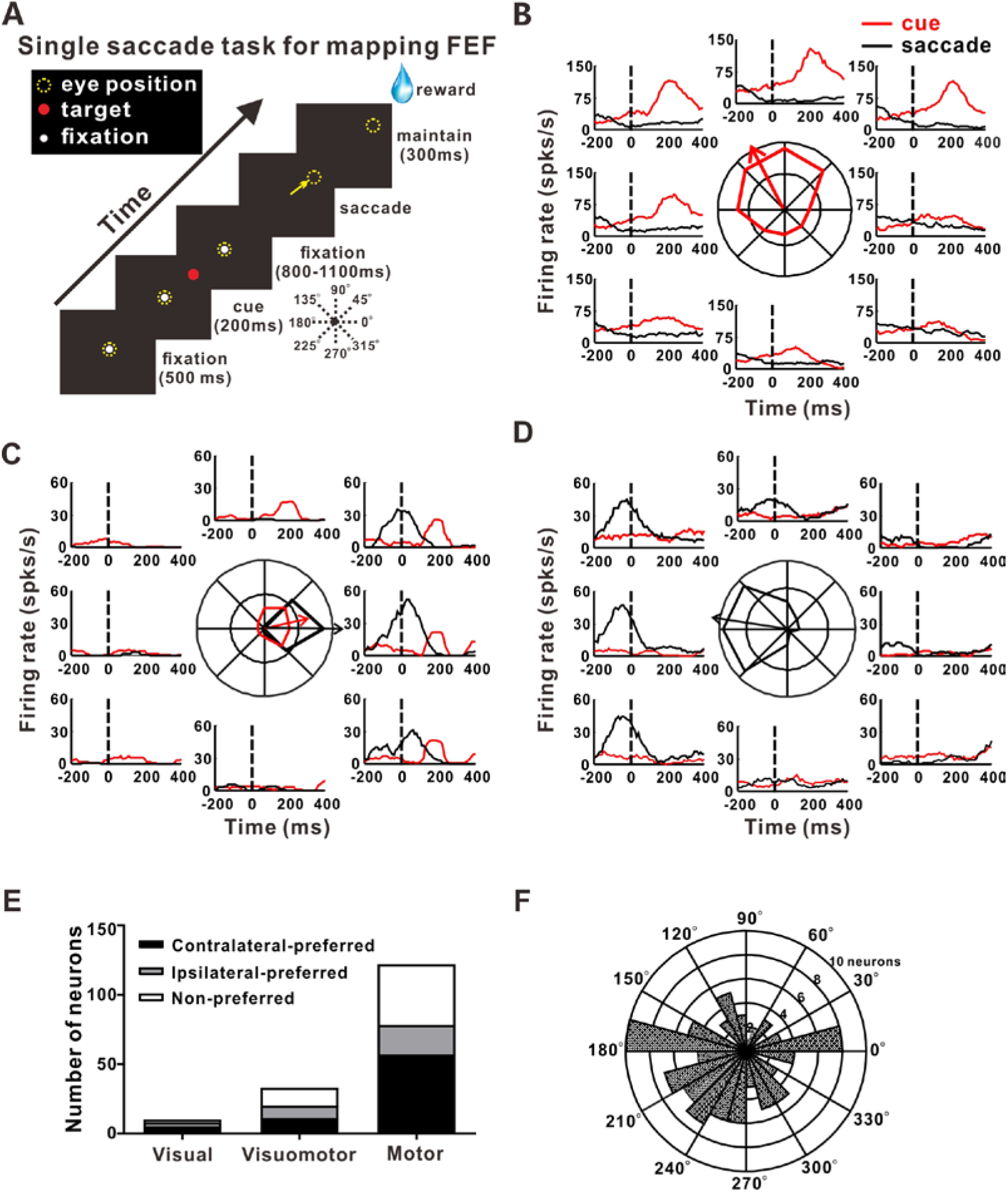
The saccadic direction tuning of FEF neurons during single memory saccade task: **(A) T**he schematic diagram of single memory-guided saccade task for mapping FEF area. In each trial, the location of targets was randomly chosen from 8 peripheral positions. **(B-D)** Representative visual and/or saccadic direction tuning plot for visual (**B**), visuomotor (**C**) and motor neurons (**D**). **(E)** Most of visuomotor and motor neurons showed significant direction preference during the single saccade (n=165 neurons, prefered: p<0.05 in Rayleigh test). **(F)** The distribution of preferred saccadic direction of visuomotor and motor neurons.

**Figure 4-1.**
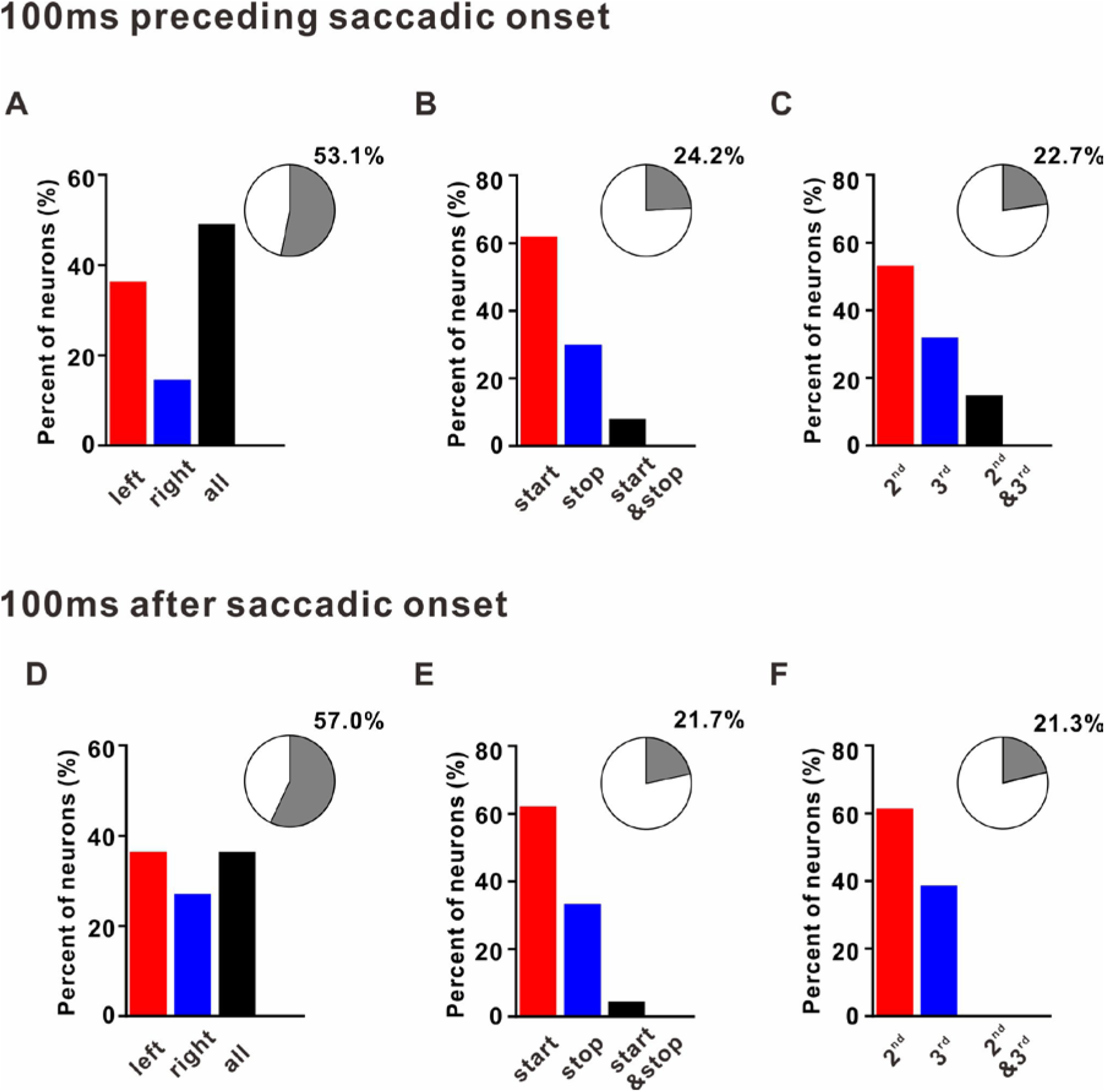
Percentages of rank activities in different analysis windows. **(A-C)** Proportion of FEF showing element **(A)**, sequence start/stop **(B)**, and subsequence switch-related **(C)** activity in pre saccade window (−100ms ~ 0). The percentage among the whole recorded FEF was presented as pie plot in the inset. **(D-E)** Proportion of FEF showing element **(D)**, sequence start/stop **(E)**, and subsequence switch-related **(F)** activity in post saccade window (0 ~100ms).

**Figure 6-1.**
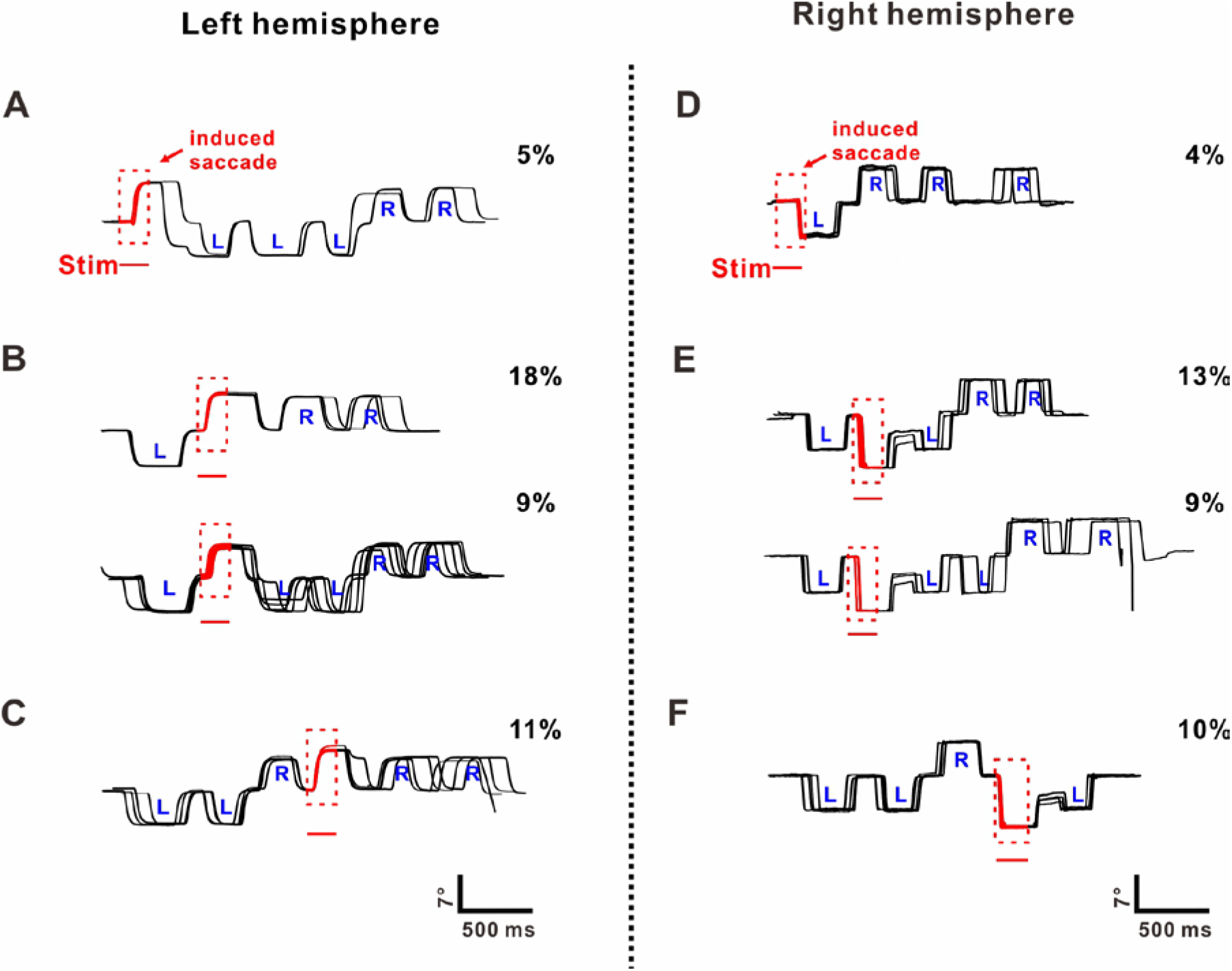
The rest of example eye traces obtained during suprathreshold microstimulation in FEF on LLRR sequential task: **(A-C)** The third main type of eye traces during suprathreshold stimulation from left hemisphere before 1^st^ saccade **(A)**, 2^nd^ saccade **(B)**, 4^th^ saccade **(C)**. Red traces represented the eye traces evoked by electrical stimulation. **(D-F)** The third and fourth main type of eye traces during suprathreshold stimulation from right FEF before 1^st^ saccade **(D)**, 2^nd^ saccade **(E)**, 4^th^ saccade **(F)**. Formats are same as the left hemisphere. All the rest percentages in (**A-B, D-E**) and some eye traces after stimulation before 3^rd^ saccade in right FEF are those trials with sequence broken after stimulation.

**Figure 6-2.**
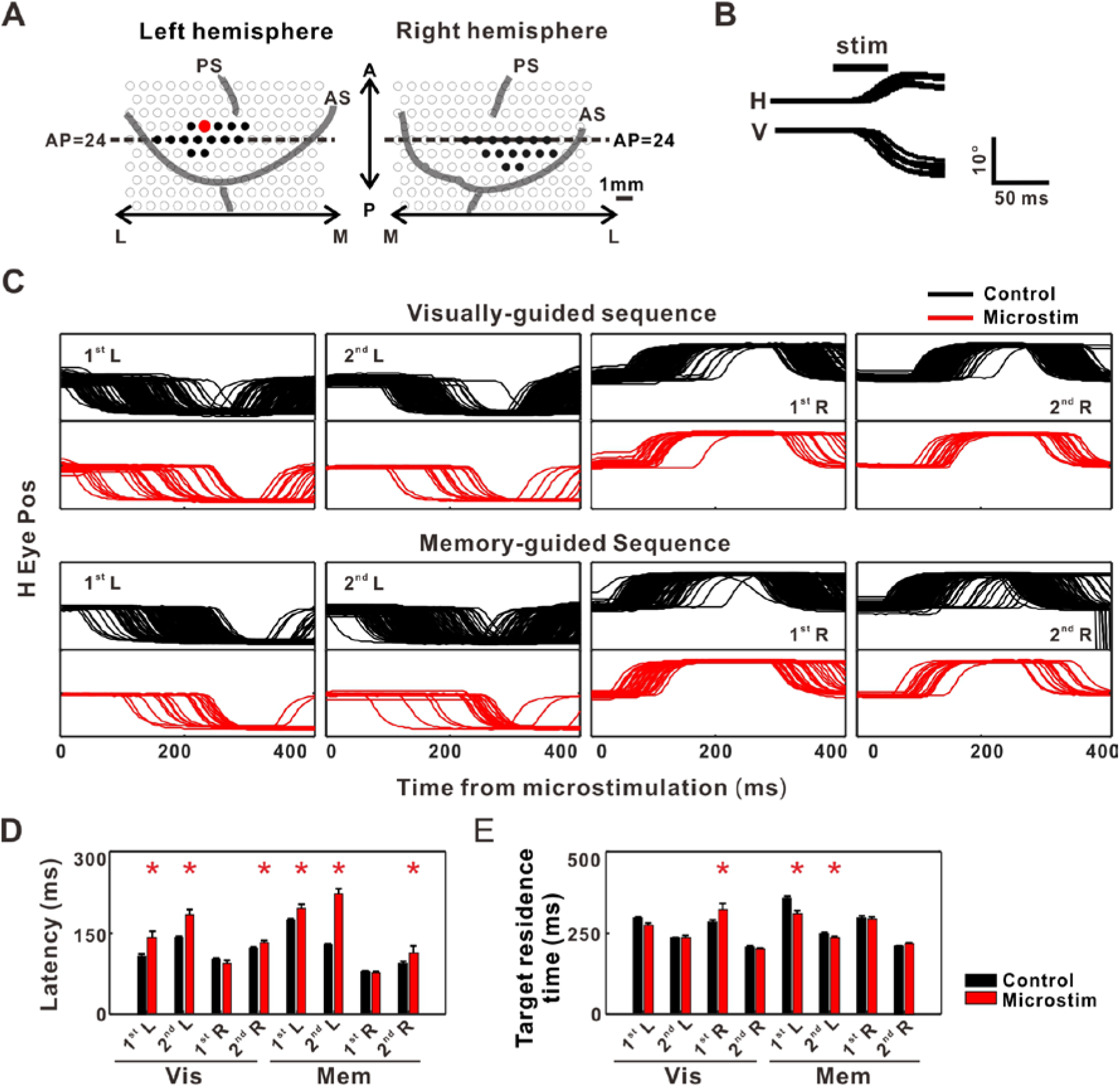
Effects of subthreshold microstimulation in FEF on sequential saccade performance. **(A)** Location of all the penetrations (black filled circles, n = 30 penetrations) and an example penetration (red filled circle) examined with subthreshold microstimulation was presented. **(B)** Eye traces of stimulation (red) and control (black) trials in one block. **(C)** Normalized ratio of response latency between stimulated and control trials. **(D)** Normalized ratio of target residence time between stimulated and control trials. For all figures, *p < 0.05, two tailed t test.

**Figure 6-3.**
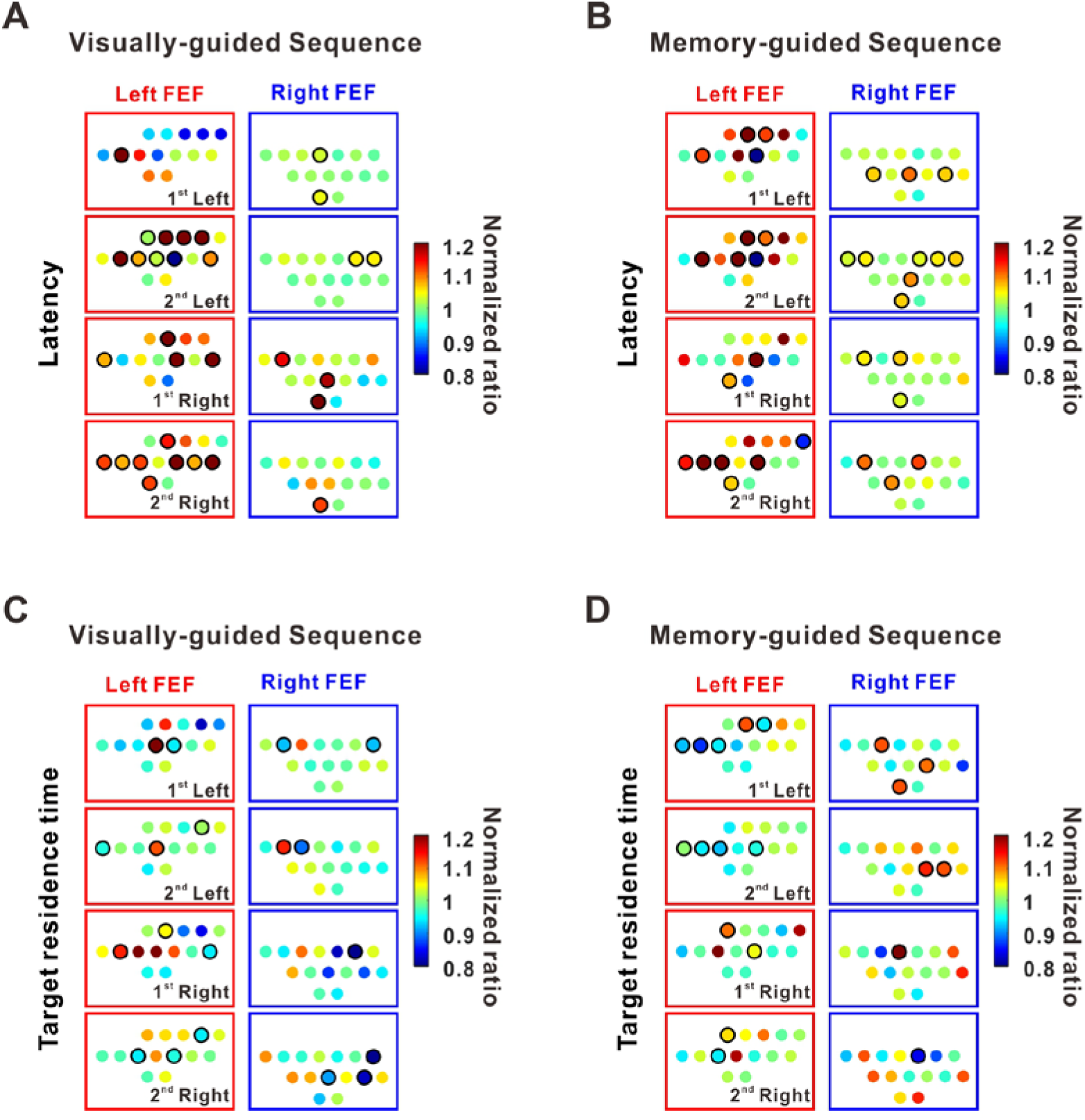
Summarized effect of subthreshold microstimulation in FEF on sequential saccade performance. **(A)** The effects on the saccade latency during visually-guided sequence, measured with a normalized ratio of saccade latency between stimulated and control trials. Each color dot represents a penetration location examined with subthreshold microstimulation (n = 30). The penetrations with significant changes in saccade latency were marked with black circles (p < 0.05, Wilcoxon signed rank test). **(B)** The effects on the saccade latency during memory-guided sequence. Formats same as **(A)**. **(C-D)** Show the effects on the target residence time during visually **(C)** and memory **(D)** guided sequential task. In general, left hemisphere was more affected by subthreshold electrical stimulation, consistent with the hypothesis that the left hemisphere is critical for sequence learning.

**Figure 8-1.**
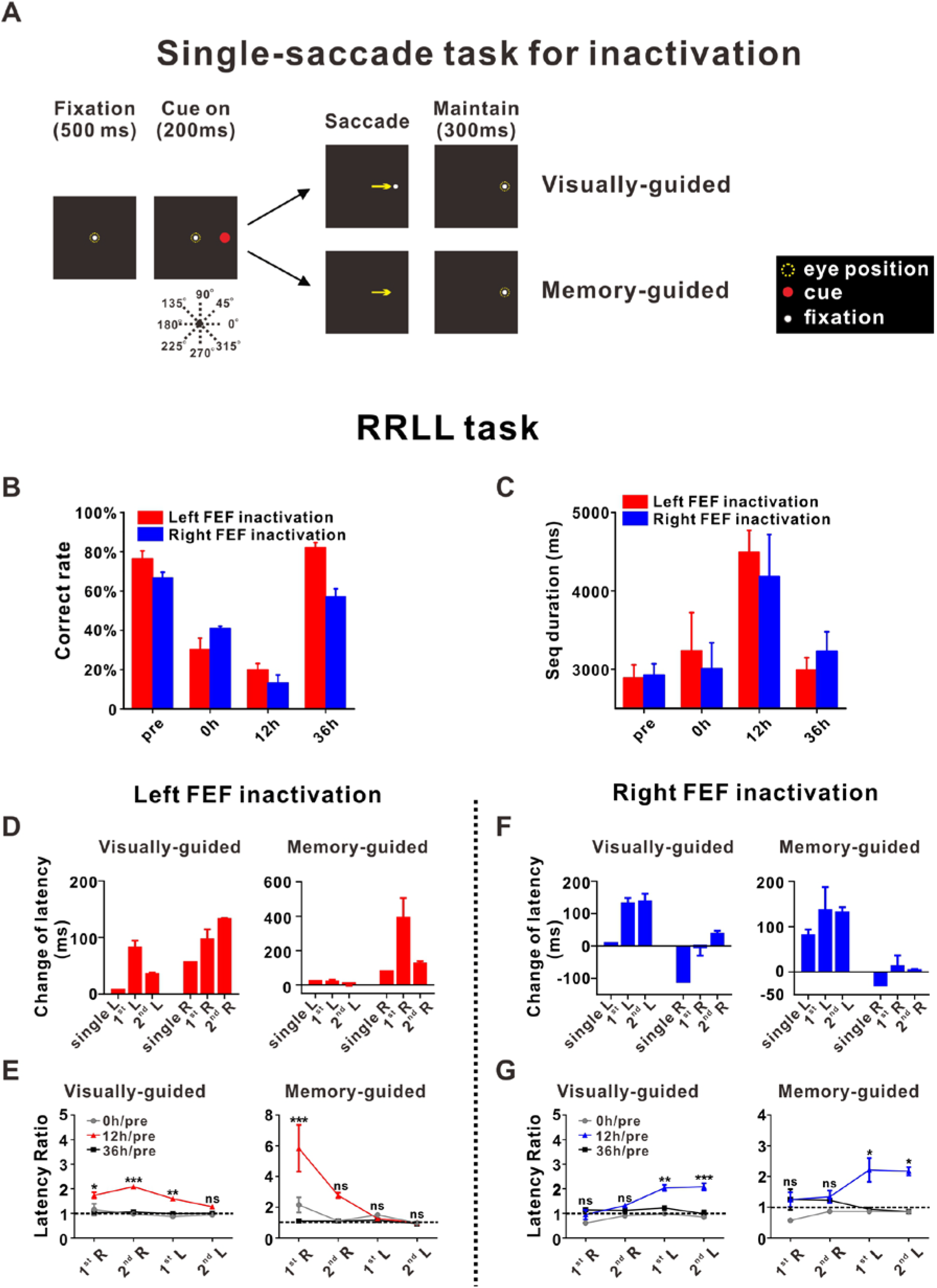
Effects of muscimol injection on RRLL sequential task. **(A)** A new visually-guided (upper panels) and memory-guide (lower panels) single saccade for inactivation experiment. A fixation spot first appeared at the center of a screen, and monkey need to maintain fixation for 500 ms. Then, a target spot appeared at either left or right periphery of the visual field for 200 ms. Once the target dispeared, for the **visually-guided single saccade**, the fixation point moved to the target position, and the monkey needs to make a saccade to the fixation point at new position, and will get reward after maintaing fixation for 300 ms; For the **memory-guided single saccade**, at the same time, the fixation spot disappeared, and the monkey needs to make a saccade to the vanished target. If the monkey did so, the target reappeared and the monkey was rewarded for continuing to fixate it for at least 300 ms. **(B)** The correct rate of RRLL sequential performance after muscimol injection in left (red) and right (blue) FEF. The abscissa marks the four different time points at which data were collected: Pre, 0, 12, or 36 h. Left hemisphere indicated byRed bars: 12h vs. pre/0h/36h, p = 2.0*10^−4^, p = 0.39 and p = 3.0*10^−4^, respectively; Right hemisphere indicated by Blue bars: 12h vs pre/0h/36h, p = 8.8*10^−7^, p = 0.026, and p = 0.0032, respectively, two tailed t test. **(C)** The effect on sequence duration at different time point after muscimol injection. Left hemisphere represented by red bars: 12h versus pre/0h/36h, p = 4.7*10^−10^, p = 0.10 and p = 2.0*10^−4^, respectively; Right hemisphere represented by blue bars: 12h vs. pre/0h/36h, p = 8.6*10^−6^, p = 0.036, and p = 0.032, respectively, two tailed t test. **(D)** Summary of latency changes at 12h time point after injection in left FEF. Visually-guided: 1^st^ L vs. 2^nd^ L, p = 0.068; 1^st^ R vs. 2^nd^ R, p = 0.29; Memory-guided: 1^st^ L vs. 2^nd^ L, p = 0.025; 1^st^ R vs 2^nd^ R, p = 0.16, one-way ANOVA with Tukey’s multiple comparisons test. **(E)** Saccade latency ratio (saccade latency after injection divided by the latency measured before injection) changes as time in left FEF. 12h vs. pre indicated by red symbols and line during visually-guided: 1^st^ R vs. 2^nd^ R, p = 0.17; 1^st^ L vs. 2^nd^ L, p = 0.22; 12h vs. pre indicated by red symbols and line during memory-guided: 1^st^ R vs 2^nd^ R, p = 0.012; 1^st^ L vs 2^nd^ L, p = 0.99, two-way ANOVA with Tukey’s multiple comparisons test. **(F)** Summary of latency changes at 12h time point after injection in righ FEF. Visually-guided: 1^st^ L vs 2^nd^ L, p = 0.97; 1^st^ R vs 2^nd^ R, p = 0.40; Memory-guided: 1^st^ L vs 2^nd^ L, p = 0.48; 1^st^ R vs 2^nd^ R, p = 0.42, one-way ANOVA with Tukey’s multiple comparisons test. **(G)** Saccade latency ratio (saccade latency after injection divided by the latency measured before injection) changes as time in right FEF. 12h vs. pre indicated by blue symbols and line during visually-guided: 1^st^ R vs. 2^nd^ R, p = 0.42; 1^st^ L vs. 2^nd^ L, p > 0.99; 12h vs. pre indicated by blue symbols and line during memory-guided: 1^st^ R vs. 2^nd^ R, p > 0.99; 1^st^ L vs. 2^nd^ L, p > 0.99, two-way ANOVA with Tukey’s multiple comparisons test. For **(E)** and **(G)**, *p < 0.05, **p < 0.001, ***p < 0.0001, and ns, no significant difference with two-way ANOVA, followed by Sidak’s multiple comparisons test. Error bars denote SEM (**B, D-G**) and STD (**C**).

**Figure 8-2.**
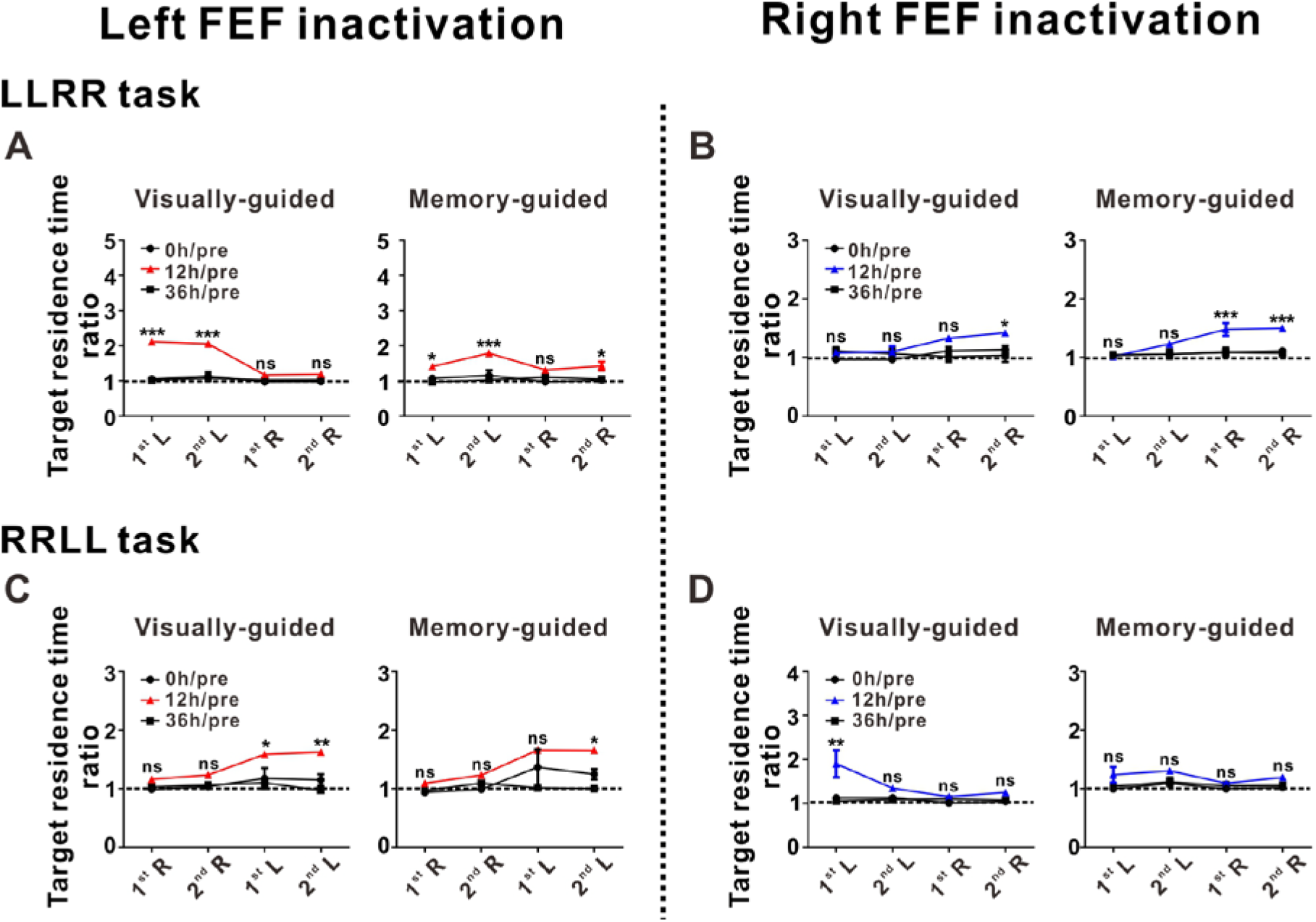
Effects of muscimol injection on target residence time for both LLRR and RRLL sequential task. **(A)** The time duration for monkey fixating at the target (measured as “Target residence time ratio” between inactivation and control trials) during LLRR sequential task after muscimol injection in left FEF. The abscissa marks the four different time points at which data were collected: pre, 0, 12, or 36 h. 12h vs. pre indicated by red symbols and line during visually-guided: 1^st^ L vs. 2^nd^ L, p = 0.89; 1^st^ R vs. 2^nd^ R, p = 0.99; 12h vs. pre indicated by red line during memory-guided: 1^st^ L vs. 2^nd^ L, p = 0.050; 1^st^ R vs. 2^nd^ R, p = 0.84, two-way ANOVA with Tukey’s multiple comparisons test. **(B)** The time duration for monkey fixating at the target (“Target residence time”) during LLRR sequential task after muscimol injection in right FEF. 12h vs. pre indicated by blue symbols and line during visually-guided: 1^st^ L vs. 2^nd^ L, p > 0.99; 1^st^ R vs. 2^nd^ R, p = 0.90; 12h vs. pre indicated by blue symbols and line during memory-guided: 1^st^ L vs. 2^nd^ L, p = 0.12; 1^st^ R vs. 2^nd^ R, p > 0.99, two-way ANOVA with Tukey’s multiple comparisons test. **(C)** The time duration for monkey fixating at the target during RRLL sequential task after muscimol injection in left FEF. Formats same as LLRR task. 12h vs. pre indicated by red line during visually-guided: 1^st^ R vs. 2^nd^ R, p = 0.93; 1^st^ L vs. 2^nd^ L, p = 0.99; 12h vs. pre indicated by red symbols and line during memory-guided: 1^st^ R vs. 2^nd^ R, p = 0.90; 1^st^ L vs. 2^nd^ L, p > 0.99, two-way ANOVA with Tukey’s multiple comparisons test. **(D)** The time duration for monkey fixating at the target during RRLL sequential task after muscimol injection in right FEF. 12h vs. pre represented by blue symbols and line during visually-guided task: 1^st^ R vs. 2^nd^ R, p = 0.024; 1^st^ L vs. 2^nd^ L, p = 0.94; 12h vs. pre represented by blue symbols and line during memory-guided sequence task: 1^st^ R vs. 2^nd^ R, p = 0.89; 1^st^ L vs. 2^nd^ L, p = 0.72, two-way ANOVA with Tukey’s multiple comparisons test. For all figures, *p < 0.05, **p < 0.001, ***p < 0.0001, and ns, no significant difference with two-way ANOVA, followed by Sidak’s multiple comparisons test. Error bars denote SEM.

## Notes

### Competing Interest Statement

The authors have declared no competing interest.

### Summary of Updates

We modified the introduction and discussion to make it more easy to understand.

